# A Circuit for Secretion-coupled Cellular Autonomy in Multicellular Eukaryotes

**DOI:** 10.1101/2021.03.18.436048

**Authors:** Lingxia Qiao, Saptarshi Sinha, Amer Ali Abd El-Hafeez, I-Chung Lo, Krishna K. Midde, Tony Ngo, Nicolas Aznar, Inmaculada Lopez-Sanchez, Vijay Gupta, Marilyn G. Farquhar, Padmini Rangamani, Pradipta Ghosh

## Abstract

Cancers represent complex autonomous systems, displaying self-sufficiency in growth signaling. Autonomous growth is fueled by a cancer cell’s ability to ‘secrete-and-sense’ growth factors: a poorly understood phenomenon. Using an integrated systems and experimental approach, here we dissect the impact of a feedback-coupled GTPase circuit within the secretory pathway that imparts secretion-coupled autonomy. The circuit is assembled when the Ras-superfamily monomeric GTPase Arf1, and the heterotrimeric GTPase Giαβγ and their corresponding GAPs and GEFs are coupled by GIV/Girdin, a protein that is known to fuel aggressive traits in diverse cancers. One forward and two key negative feedback loops within the circuit create closed-loop control (CLC), allow the two GTPases to coregulate each other, and convert the expected switch-like behavior of Arf1-dependent secretion into an unexpected dose response alignment behavior of sensing and secretion. Such behavior translates into cell survival that is self-sustained by stimulus-proportionate secretion. Proteomic studies and protein-protein interaction network analyses pinpoint growth factors (e.g., the epidermal growth factor; EGF) as a key stimuli for such self-sustenance. Findings highlight how enhanced coupling of two biological switches in cancer cells is critical for multiscale feedback control to achieve secretion-coupled autonomy of growth factors.

**SYNOPSIS IMAGE:** 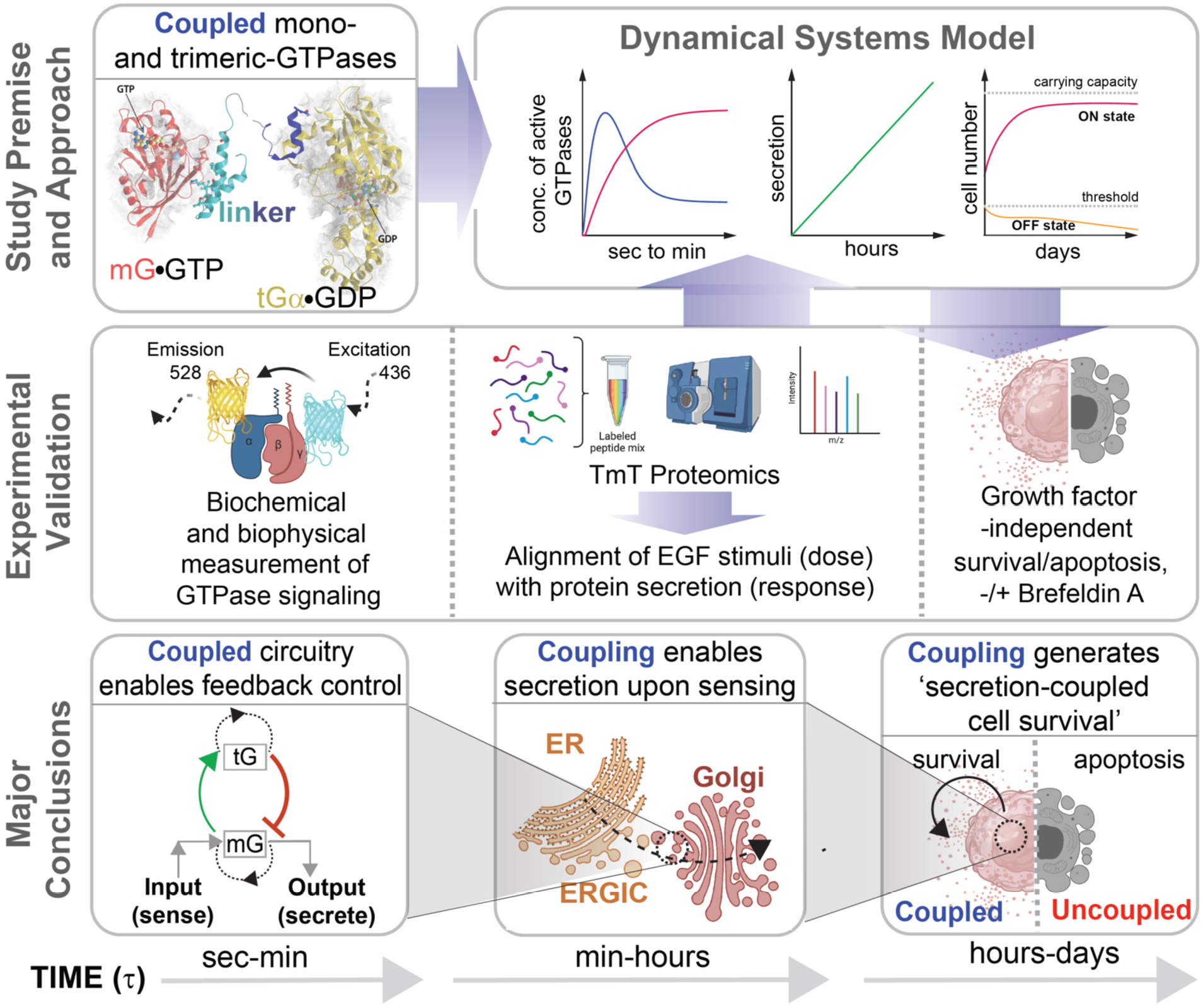

**STANDFIRST TEXT:** This work defines the inner workings of a Golgi-localized molecular circuitry comprised of coupled GTPases, which empowers cells to achieve self-sufficiency in growth factor signaling by creating a secrete-and-sense autocrine loop.

**HIGHLIGHTS/MAIN FINDINGS:**

- Modeling and experimental approaches were used to dissect a coupled GTPase circuit.
- Coupling enables closed loop feedback and mutual control of GTPases.
- Coupling generates dose response alignment behavior of sensing and secretion of growth factors.
- Coupling is critical for multiscale feedback control to achieve secretion-coupled autonomy.

## INTRODUCTION

Self-sufficiency in growth signaling, a.k.a, growth signaling autonomy is the first of the six hallmarks of all cancers to be clearly defined (Hanahan and Weinberg, 2000). While most growth factors (GFs) are made by one cell type in order to stimulate proliferation of another, many cancer cells synthesize GFs to which they are responsive, creating a positive feedback signaling loop called autocrine stimulation (Fedi P, 1997). Serum-free cell culture studies squarely implicate such stimulation as key support for intracellular mechanisms that impart autonomy (reviewed in (Chigira et al., 1990)). Autonomy in cancer cells obviates dependence on extrinsic GFs, as illustrated in the case of PDGF (platelet-derived growth factor) and TGFα (tumor growth factor α) in glioblastomas and sarcomas, respectively (Fedi P, 1997). Beyond cancers, ‘secrete-and-sense’ circuits that allow cells to secrete and sense the same signaling molecule are ubiquitous (Youk and Lim, 2014); these autocrine secrete-and-sense mechanisms don’t just enable autonomy (Maire and Youk, 2015) but also generate diverse social behaviors, and recur across species (Youk and Lim, 2014).

Autocrine secretion of growth factors relies on an essential, efficient, and accurate molecular machinery that constitutes a central paradigm of modern cell biology, i.e., the secretory pathway (Matlin and Caplan, 2017; Trombetta and Parodi, 2003). This pathway consists of various modules that are compartmentalized on the endoplasmic reticulum (ER) and the Golgi apparatus, and are responsible for folding, processing of the post-translational modifications (PTMs), and trafficking of the proteins routed to the membrane of extracellular space (Kelly, 1985; Rothman and Orci, 1992). Nearly all these aspects of the secretory pathway have been found to be deregulated in cancers, ranging from observed changes in Golgi shape (‘onco-Golgi’ (Petrosyan, 2015)), or its function (Zhang, 2021), which inspired the development of disruptors of this ER-Golgi secretory system as anti-cancer agents (Luchsinger et al., 2018; Núñez-Olvera et al., 2020; Ohashi et al., 2012; Ohashi et al., 2016; Ohashi et al., 2017; Wlodkowic et al., 2009).

Despite these insights, the core mechanisms of cell secretion that impart cell autonomy remains poorly understood. To begin with, it is still unknown whether or not secretion is proportional to growth factor stimulation, and whether such secretion is sufficient to support cell survival, perhaps via closed loop autocrine sensing and signaling (the so-called ‘secrete-and-sense’ loop (Youk and Lim, 2014)). A recent study has shown that the secretory functions of the Golgi apparatus requires the unlikely coupling of two distinct species of GTPases at the Golgi (Lo et al., 2015) (**Figure 1A**): one is small or monomeric (m) GTPase Arf1, and the other is heterotrimeric (t) GTPases Gi. GTPases serve as molecular switches that gate signal transduction: “on” when GTP-bound (active) and “off” when GDP-bound (inactive). The “ADP-ribosylation factor” (Arf1) (Kahn and Gilman, 1986) mGTPase is localized to the Golgi complex in mammalian cells and is essential for the secretory pathway (Stearns et al., 1990); it associates with Golgi membranes upon activation, and is released from Golgi membranes into the cytosol upon inactivation. Such cycles of association and dissociation are regulated by Golgi-associated, guanine nucleotide exchange factors (GEFs) and GTPase activating proteins (GAPs). Trimeric GTPases were detected in the Golgi over three decades ago (Barr et al., 1992; Stow et al., 1991), and numerous studies have provided clues that they may regulate membrane traffic and maintain the structural integrity of the Golgi (reviewed in (Cancino and Luini, 2013)). However, the concept of G protein activation at the Golgi and the potential impact of such activation remained controversial, primarily due to the lack of direct proof of G protein activation. The study that reported coupling of Arf1 mGTPase and Giαβγ tGTPase provided direct evidence, the first of its kind, that the two GTPases are coupled by a linker protein, Gα-Interacting Vesicle-associated protein (GIV) (Lo et al., 2015). Activation of Arf1 mGTPase facilitates the recruitment of GIV on the membrane via a direct, nucleotide-dependent interaction. Upon recruitment, GIV binds and activates Gαi serving its role as a GEF for the tGTPase, Gi. Such activation of Gi at the Golgi affects two fundamental functions of the Golgi, i.e., vesicle trafficking and the structural organization of the Golgi stacks--both via modulation of Arf1 signaling. These findings firmly established that Gαi is functionally active in the Golgi.

**Figure 1.**
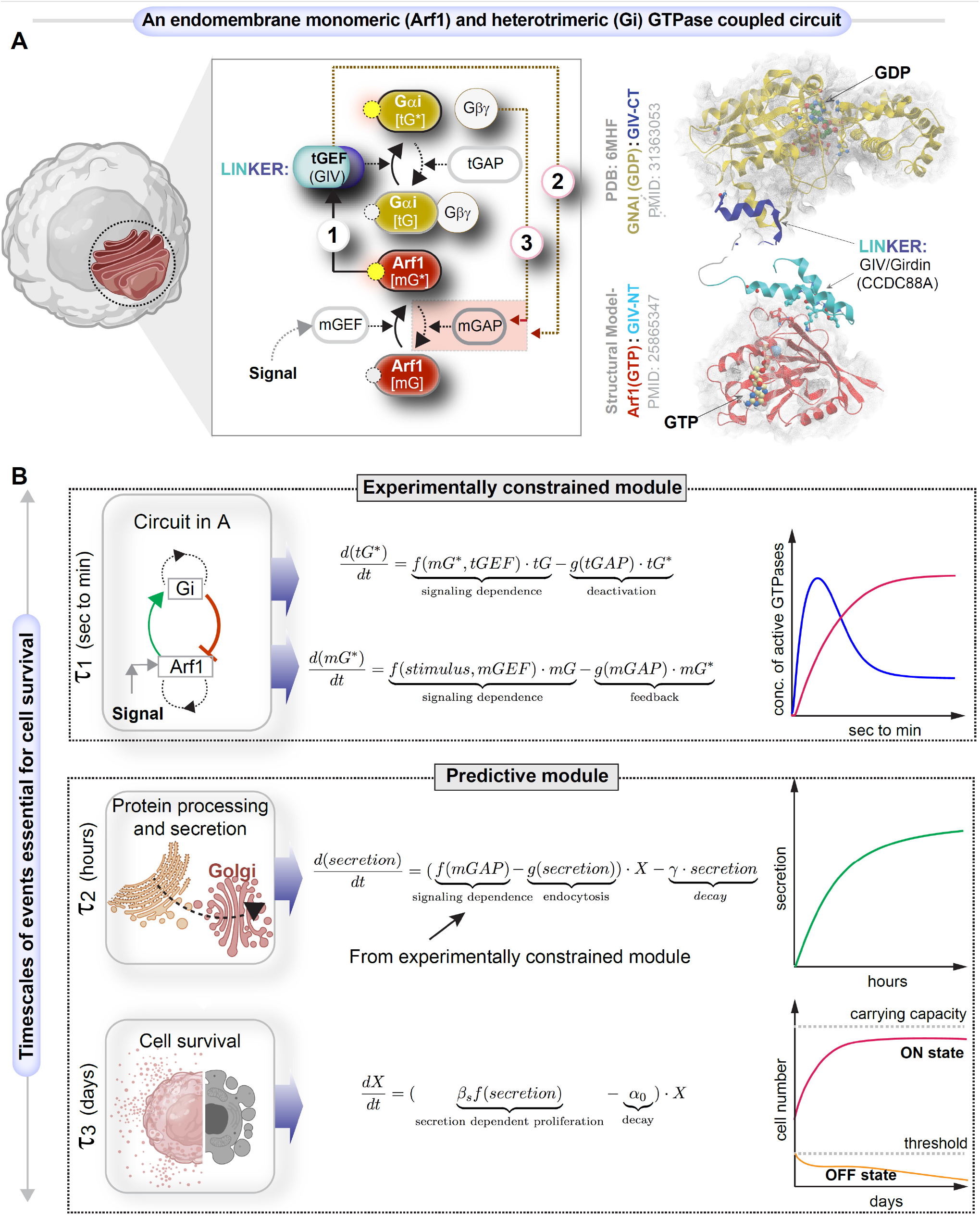
Study design and approach. **A.** Schematic shows a system of two species of GTPases, mGTPases (mG) and heterotrimeric GTPases (tG), coupled by a linker protein, GIV/Girdin, that is localized on the Golgi membranes within the secretory pathway as the focus of this study. The circuit begins when active Arf1-GTP directly binds GIV’s N-term HOOK domain, recruits GIV to Golgi membranes, and activates Gi (Lo et al., 2015) (arrow 1). The circuit is completed when GIV’s C-terminus orchestrates two feedback loops (arrows 2 and 3), both of which are essential for the inactivation of Arf1 (Kalogriopoulos et al., 2019; Lo et al., 2015). See also **Figure EV1** for illustrations detailing the sequential steps within the dynamic nature of the motif, and **Movie EV1** for the visualization of these dynamic steps as a movie gif. **B**. Schematic of the dynamical systems model that we used to study the role of such coupling of GTPase (top panel) in autocrine secretion-supported cell survival and proliferation (bottom panel). The modeling in the top is experimentally constrained, and the modeling in the bottom is a predictive module. This model is based on the nominal time scale of these events (left panel) and has the typical behavior shown in the right panel.

Because tGTPases are known to primarily transduce extracellular signals (‘sensing’) into intracellular signals that shape cellular responses, we asked how coupling of the two GTPases, one that guards cell secretion (Arf1) and another that gates signal sensing (Gi), may impact the cell’s ability to secrete-and-sense. In systematically interrogating this question, we viewed the experimentally validated interactions and functions of the two GTPases and their GEFs and GAPs as a circuit of coupled GTPases. Such coupling, whose structural basis has been experimentally validated (**Figure *1A-right***), forms a closed loop that is comprised of one forward reaction and two negative feedback loops (**Figure *1A-left;* Figure EV1; Movie EV1; Materials and Methods**). The forward reaction is the recruitment of GIV/Girdin by active Arf1 on Golgi membranes (arrow 1). GIV is a multi-modular cytosolic signal transducer that is a prototypical member of the family of guanine nucleotide exchange modulators (GEM) of tGTPases; GIV’s GEM domain binds and activates the tGTPase Gαi, and thereby, serves as a tGEF within this circuit. One negative feedback loop is that GIV can improve the GAP for Arf1---ArfGAP2/3, thus terminating Arf1 signaling (arrow 2); the other is due to GIV’s role as GEF to activate Gi and thus enhance Arf1GAP2//3, which also lead to the termination of Arf1 signaling (arrow 3). This phenomenon of co-regulation between the two classes of GTPases maintains Golgi shape and function, two closely intertwined processes that are regulated by Arf1. The triggers for and the consequence(s) of such co-regulation on signal sensing/response remained unknown.

Because coupling of two species of GTPase switches, Arf1 and Gi, with feedback control is likely to generate complex, nonlinear, and non-intuitive emergent properties, we use cross-disciplinary approaches to dissect the role of the coupled GTPases within the secretory pathway and explore its functional significance in eukaryotic cells. Using systems biology approaches and explicit integration of experimental biology and computational methods, we also assess the impact of perturbing this motif, i.e., uncoupling the GTPases. Our findings show how coupling makes secretion responsive to growth factors, in particular the epidermal growth factor (EGF), and appears to impart secretion-coupled autonomy.

## RESULTS

### An integrated systems and experimental approach to dissect a Golgi-localized GTPase circuit

We began by developing a dynamical systems model for this coupled circuit (**Figure 1B**; see **Materials and Methods**) and drawing clues from protein-protein interaction (PPI) network analyses, to generate testable hypotheses and validate them experimentally. The integrated approach allowed us to connect across time scales of the emergent behavior of the coupled GTPase circuit with cellular secretion, cell survival, and ultimately, secretion-coupled survival, i.e., autocrine autonomy.

The first part of the dynamical systems model is an experimentally constrained module for the coupled GTPases switches (upper panel in **Figure 1B**), where normalized Hill functions are used (Cao et al., 2020; Saucerman and McCulloch, 2004) (see **Materials and Methods** for details). This approach was chosen to capture the key timescales and molecular players involved rather than focus on the specific biochemical reactions. Additionally, this approach has fewer free parameters than the traditional approach of building networks with large numbers of reactions (Getz et al., 2019), leading to less ambiguity of decision making for model development. The kinetic parameters of the coupled GTPases module were subsequently tuned to fit the time course data of GTPases in control cells and GIV-depleted cells.

The second part of the dynamical systems model is a predictive module for cell secretion and secretion-coupled cell survival (lower panel in **Figure 1B**). The coupling of this predictive module with the above experimentally constrained module is achieved by setting the secretion rate as a function of mGAP. The following findings allow us to make this coupling in the model: the finiteness of Arf1 activation-inactivation cycle was assumed to be a surrogate indicator of successful anterograde cargo movement through the compartments within the secretory pathway, i.e., the ERGIC (endoplasmic-reticulum–Golgi intermediate compartment) to the Golgi, because Arf1 regulates membrane traffic through a cycle of GTP binding and hydrolysis (Donaldson and Jackson, 2011); GTP binding is a pre-requisite for membrane curvature and vesicle formation (Beck et al., 2008) from the donor compartment, whereas GTP hydrolysis is a pre-requisite for vesicle uncoating (Tanigawa et al., 1993) and fusion with acceptor compartment. Therefore, we set the secretion rate as a function of GTP hydrolysis, a process regulated by mGAP. Except this setting for secretion rate, the model for cell secretion and cell survival/proliferation is similar to the model proposed by Hart *et al* (Hart et al., 2014), where the kinetic parameters are from biologically plausible ranges reported previously (Adler et al., 2018).

Two different cancer cell lines were chosen to conduct experiments, the HeLa cervical cancer and the MDA-MB231 breast cancer cells. Our choice was guided by two reasons: (*i*) HeLa cells not only represent the most robust system to study Golgi structure (Ayala and Colanzi, 2016; Wortzel et al., 2017) and function (Rauter et al., 2020), but also provide continuity with prior work because all biophysical and functional studies that led to the discovery of the coupled GTPases at the Golgi were performed in this model; (*ii*) we and others have shown that transcriptional upregulation or post-transcriptional activation (Bhandari et al., 2015; Dunkel et al., 2012; Sasaki et al., 2015) of GIV (the ‘linker’ between the two GTPases; **Figure 1A**) supports several aggressive tumor cell properties (of which, many were demonstrated in MDA-MB231 cells (Jiang et al., 2008; Lopez-Sanchez et al., 2015; Midde et al., 2018; Rahman-Zaman et al., 2018; Rohena et al., 2020; Wang et al., 2015; Wang et al., 2017)), including, invasion, matrix degradation, proliferation and survival (Aznar et al., 2016; Garcia-Marcos et al., 2015). Elevated expression of GIV has also been reported in a variety of solid tumors (Garcia-Marcos et al., 2015; Getz et al., 2019), both in primary tumors (Ghosh, 2015; Ghosh et al., 2016b) as well as in circulating tumor cells (Barbazan et al., 2016; Dunkel et al., 2018) have been shown to correlate with tumor aggressiveness and poor survival across cancers. Finally, model and PPI network-driven predictions of uncoupling the GTPases or interrupting secrete-and-sense autonomy were experimentally validated in the two cancer cell lines that lack GTPase coupling in the absence of the GIV linker protein.

### EGF activates Arf1 (mG*) at the Golgi and triggers the recruitment of a GEF for trimeric Giαβγ

First, we sought to model the impact of coupling on m/tGTPase signaling in response to input signal (**Figure 1A**). Key events within the circuit were measured experimentally using available tools and experimental approaches (Arrows 1-3; **Materials and Methods**). Epidermal growth factor (EGF) was prioritized as input signal because of prior evidence documenting its role in the regulation of Golgi secretion (Blagoveshchenskaya et al., 2008), its fragmentation during mitosis (Shaul and Seger, 2006), and most importantly, in the activation of Arf1 (Boulay et al., 2008; Haines et al., 2014; Haines et al., 2015).

We measured Arf1 activity in response to EGF using an established pull-down assay (**Figure 2A-B**). with the Glutathione S Transferase (GST)-tagged GAT domain of GGA3; this domain is known to selectively bind the active GTP-bound pool of Arf1 (Cohen and Donaldson, 2010). The levels of Arf1•GTP were increased ~3-fold within 5 min after ligand stimulation, followed by a return towards baseline by 30 min, which we assume reflects the level of Arf1 activity in cells at a steady-state (**Figure 2B**). These temporal dynamics were used to fit the parameters for Arf1 activity in the computational model of the circuit (blue line in **Figure 2C**) (R^2^ and normalized RMSE are 0.72 and 0.19 respectively; see **Materials and Methods** and **Table EV1** for model parameters). Such fitting completed the characterization of the first GTPase switch, i.e., Arf1; in this case, the input is ligand stimulus (EGF) and the output is Arf1-GTP (OUTPUT #1; mG*).

**Figure 2.**
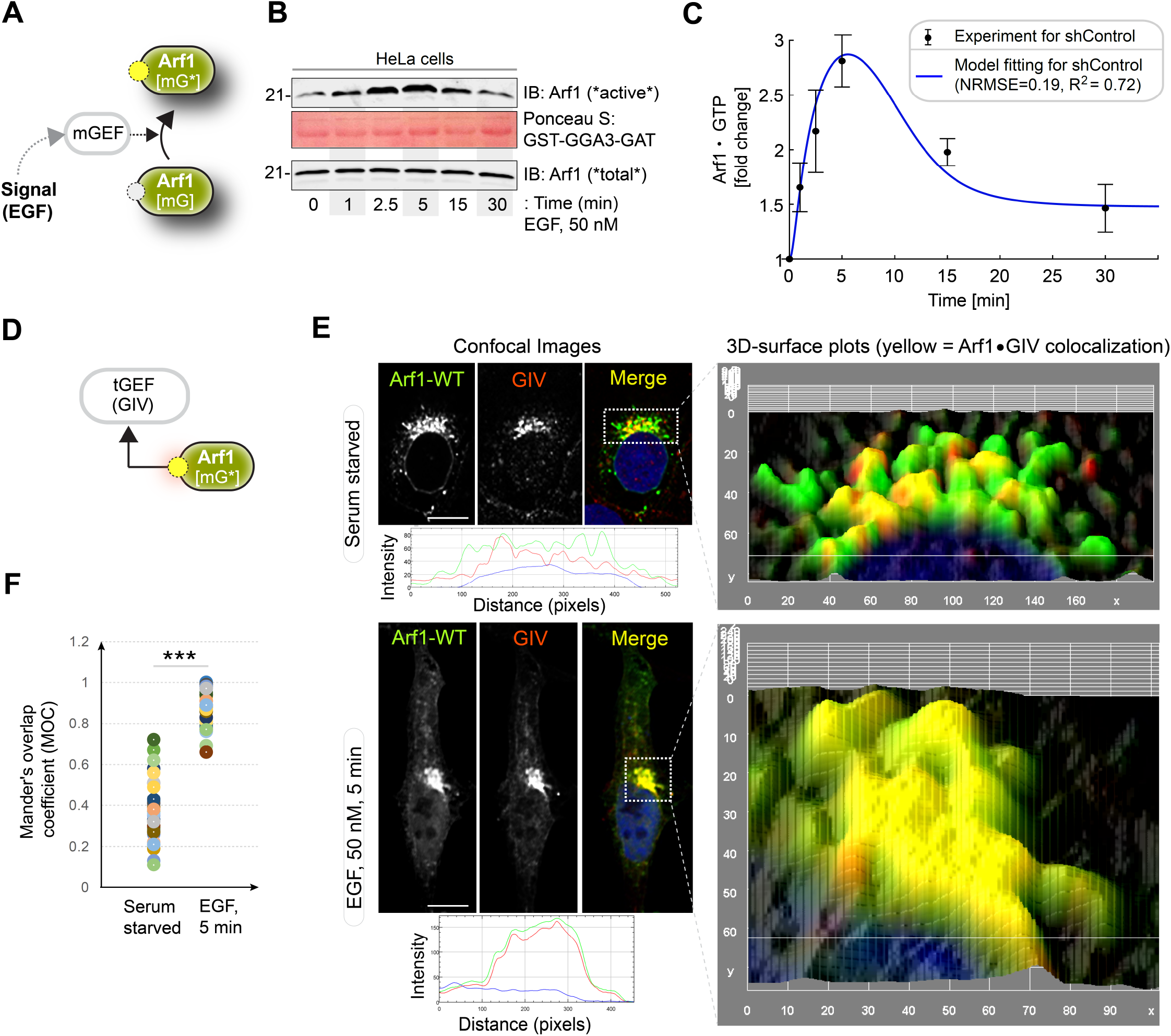
EGF activates Arf1 (mG*) within the Golgi-localized endomembrane GTPase system, triggers the recruitment of GIV-GEM on Golgi. **A**. Schematic showing the specific step being interrogated in panels B-D, i.e., Arf1 activation under EGF stimulation. **B.** Immunoblot shows GST-GGA-GAT domain bound Arf1 (active; top) and total Arf1 (input lysates; bottom) from equal aliquots of lysates of HeLa cells that were stimulated with EGF for the indicated time points prior to lysis. **C.** Graphs display the model fitting for Arf1 activation dynamics. The experimentally determined Arf1 activation (in B) dynamics are displayed as black dots with error bars, representing mean ± S.E.M (n=3), and numerical simulation is shown by the blue continuous line. **D**. Schematic showing the specific step being interrogated in panels F-G, i.e., recruitment of GIV-GEM on Golgi. **E**. HeLa cells expressing Arf1-HA were serum starved overnight (E, top) and subsequently stimulated with EGF for 5 min (E, bottom) prior to fixation with PFA. Fixed cells were stained for Arf1 (HA; green) and GIV (red) and nuclei (DAPI; blue). Panels on the left show overlay of all 3 stains and representative RGB plots of sections through the Arf1-stained pixels. Panels on the right display the magnified 3D surface plots of the boxed regions in the left panels. Scale bar = 10 μm. **F.** Scatter plot shows the Mandler’s overlap coefficient (MOC) for Arf1-HA and GIV colocalization in E that was calculated on 13-15 cells/experiment, n = 3 independent experiments. *p* values were determined using Mann-Whitney t-test: ***, 0.0002.

A key consequence of Arf1 activity within the coupled GTPase circuit is the first segment of the Gi activation pathway, i.e., the recruitment of GIV (**Figure 2D**), which is not only an effector of Arf1 but also the GEF of Gi (Lo et al., 2015). Previous studies showed that an evolutionarily conserved region in the N-terminal Hook domain of GIV can directly and preferentially bind to the active GTP-bound conformation of Arf1 (Lo et al., 2015), revealing the structure basis of the recruitment of GIV by active Arf1 (**Figure 1A**-*right*). To test whether GIV recruitment occurs in cells responding to EGF, we used immunofluorescence microscopy to observe HA-tagged Arf1 (green; **Figure 2E**) and endogenous GIV (red; **Figure 2E**). Membrane-colocalization of Arf1 and GIV was significantly increased within 5 min after EGF stimulation for serum-starved cells, as determined by a quantification of the Arf1-positive Golgi regions using a Mander’s overlap coefficient (MOC) (**Figure 2F**). These results indicate that EGF-induced Arf1 activity triggers the recruitment of GIV at the Golgi.

### EGF triggers the activation of Gi (tG*) on Golgi membranes, and then activates ArfGAP, terminating Arf1 signaling *via* feedback loops within the closed loop system

We next evaluated the second segment of the Gi activation pathway, i.e., the ability of membrane-recruited GIV to bind and activate the tGTPase Gi at the Golgi (**Figure 3A**). To be more specific, we compared Gi activation level between control cells and GIV-depleted cells. The Gi activation level is measured by a well-established FRET-based assay (Gibson and Gilman, 2006). In this assay, the αi and βγ subunits of Gi were tagged with YFP and CFP, respectively; if Gi is activated, i.e., the α and βγ subunits dissociate, YFP and CFP stay far from each other, leading to low FRET (**Figure 3B**; See **Materials and Methods** for details). Besides, the GIV-depleted cells were obtained using short hairpin RNA to target GIV (shGIV cells; **Figure 3C**; See **Materials and Methods** for details). When we conducted FRET assays in these cells, we found that there was a significant drop in FRET (i.e., activation of Gi and trimer dissociation after EGF stimulation) at the Golgi within 5 min after EGF stimulation in control cells. Activation of Gi continued to peak by 15 min in control cells, reaching a plateau by 25-30 min (**Figure 3D** *top;* **Figure 3E**; See **Figure EV2A-B** for FRET at the PM). We noted that the temporal propagation of the input signal (EGF) takes ~5 min to trigger events at the Golgi, which is considerably delayed compared to most of the well-defined EGF-stimulated, receptor-proximal events (**Figure EV4A**) which begin within ~2-5 sec (Reddy et al., 2016). This delay is consistent with the concept of propagation delay in networks (Brent, 2009). On the other hand, in shGIV cells, such tGTPase activation was abolished due to the non-changed FRET (**Figure 3D** *bottom;* **Figure 3E**). Taken together, these results demonstrate that Gi is activated at the Golgi upon EGF stimulation, and that such activation requires GIV.

**Figure 3.**
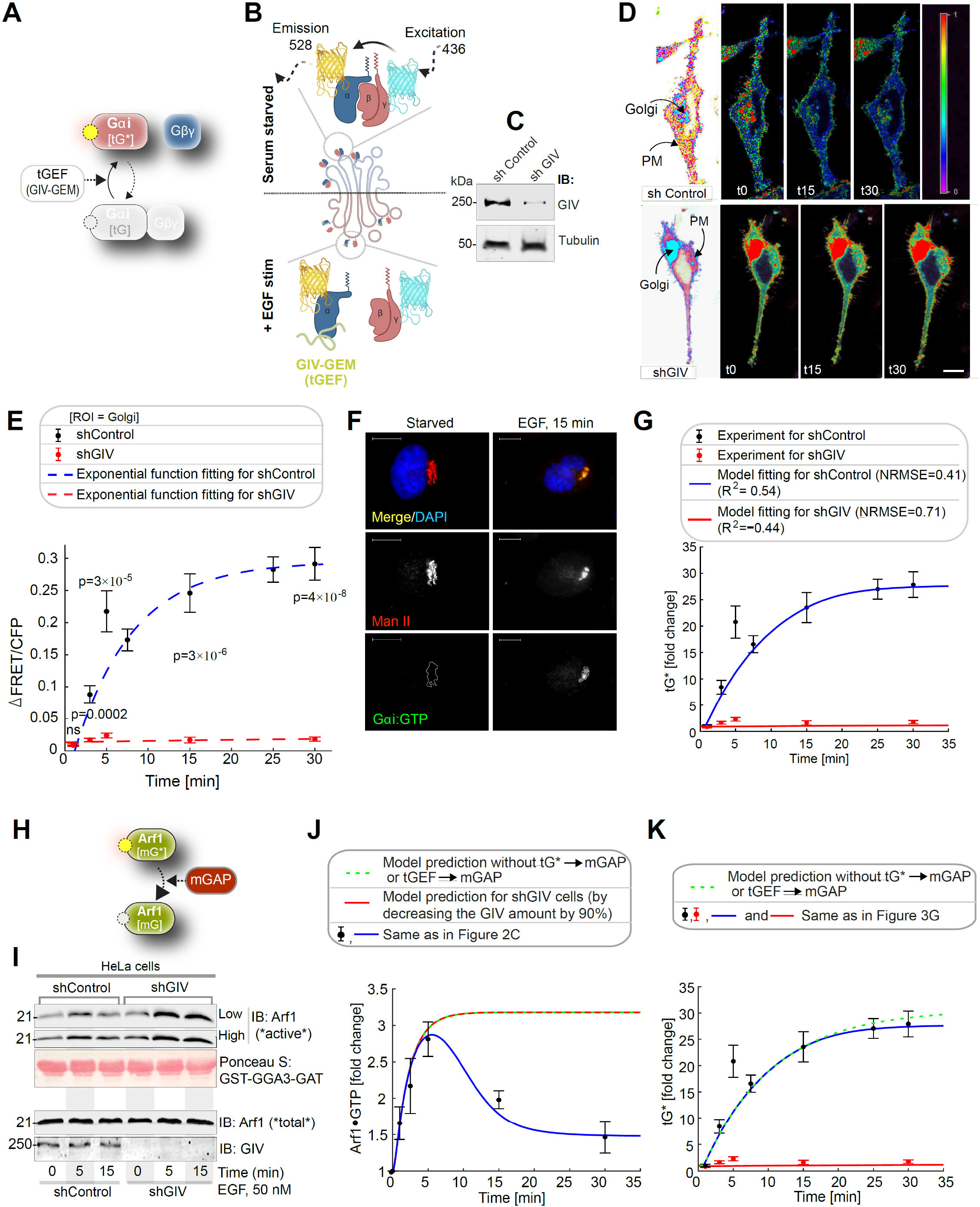
EGF triggers activation of Gi (tG*) on Golgi membranes, and then activates ArfGAP, terminating Arf1 signaling *via* feedback loops within the closed loop system. **A.** Schematic showing the specific step being interrogated in this figure, i.e., Gi activation. **B**. Schematic describing the mechanism of the FRET Gαi activity reporter. Serum starved conditions are expected to have more inactive trimeric Gi, and hence show high FRET (top). Upon ligand stimulation GIV-dependent dissociation of trimers is expected, with a resultant loss of intermolecular FRET. **C**. Equal aliquots (~45 μg) of whole cell lysates of control (shControl; top) and GIV-GEM depleted (shGIV; bottom) HeLa cells were analyzed for GIV and tubulin (loading control) by immunoblotting (IB). **D**. Control (sh Control; top) and GIV-GEM depleted (shGIV; bottom) HeLa cells were co-transfected with Gαi1-YFP, Gβ1-CFP and Gγ2 (untagged) and live cells were analyzed by FRET imaging at steady-state, after being serum starved in 0.2% FBS overnight and then after stimulation with 50 nM EGF. Representative freeze-frame FRET images are shown. FRET image panels display intensities of acceptor emission due to efficient energy transfer in each pixel. FRET scale is shown in the inset. Golgi and PM regions of interest are indicated with arrows. Scale bar = 10 μm. See also **Figure EV2A-B** for free-frame images for additional time points in control HeLa cells. **E.** ΔFRET/CFP at the Golgi (derived from D) as a function of time. The data is represented as mean ± S.E.M. Interrupted lines display the fitting results by using exponential functions for shControl (blue) and shGIV cells (red). Data represent 5 regions of interest (ROIs) analyzed over the pixels corresponding to the Golgi of 3-5 cells from 2 independent experiments. *p* values, as determined against t0 using Mann-Whitney are displayed. **F**. HeLa cells starved with 0.2% FBS overnight or stimulated subsequently with 50 nM EGF were fixed and stained for active Gαi (green; anti-Gαi:GTP mAb) and Man II (red) and analyzed by confocal microscopy. Activation of Gαi was detected exclusively after EGF stimulation. When detected, active Gαi colocalizes with Man II (yellow pixels in merge panel). See also **Figure EV2C-D** for additional time points and stimulus. Scale bar = 7.5 μm. **G.** Model fit for the fold change of active tGTPase (denoted as tG*). Experiment data is the fold change of ΔFRET/CFP in D. Continuous lines display the model simulation results after parameter fitting (See **Table EV1** for parameters). **H.** Schematic shows the step being interrogated in this figure, i.e., the termination of Arf1 signaling. **I.** Immunoblot shows bound Arf1 (active; top) and total Arf1 (input lysates; bottom) from equal aliquots of lysates of control (sh Control) and GIV-depleted (shGIV) HeLa cells. Cells were stimulated with EGF for the indicated time points prior to lysis. Bar graphs in **Figure EV2E** display the fold change in Arf1 activity normalized to t0 min. ‘Low’ and ‘high’ indicate exposures. **J-K.** Model predictions of Arf1 activation dynamics (J) and Gαi activation dynamics (K) when negative feedbacks do not exist. The depletion of negative feedbacks in the model is achieved by deleting either *tG* → mGAP* (interrupted yellow line) or *tGEF → mGAP* (interrupted green line). These two depletion ways have no difference due to AND gate logic; please see also **Figure EV3** for model predictions using OR logic. The red line in J was obtained by setting the GIV amount to 10% of the control cell, matching the low concentration of GIV in shGIV cells. As a reference, results of the simulation fit to experimental data in **Figure 2C** and **Figure 3G** are also displayed here.

Because FRET studies require the overexpression of G protein subunits at levels much higher than relevant in physiology, we sought to validate our FRET-based findings on endogenous Gi. To this end, we performed confocal immunofluorescence microscopy using a *bona fide* marker of the organelle, the Golgi-localized α-mannosidase II (Man II) (Zuber et al., 2000), and anti-Gαi•GTP mAb, which selectively recognizes the active (GTP-bound) conformation of the G protein (Lane et al., 2008). These signals colocalized not only in EGF-stimulated cells (**Figure 3F**) but also in cells exposed to other stimuli, for example, 10% serum (representing a well-mixed growth factors) and lysophosphatidic acid (LPA), a ligand for the GPCR, LPA-receptor (LPAR; **Figure EV2C-D**). Findings confirmed that Gi is activated on Golgi membranes after growth factor stimulation and suggested the prevalence of this event in response to diverse stimuli.

We fit the above experiment data by tuning the kinetic parameters. We obtained a good fit for the fold change of Gi activation in both control and shGIV cells (**Figure 3G;** R^2^ and RMSE, 0.54 and 0.41 for control cells; −0.44 and 0.71 for shGIV cells). The low level of GIV in shGIV cells was mimicked by decreasing the levels of expression of GIV to 10% of that in control cells (**Figure 3C**). Thus, the model matched the overall trend of experiment data in both cells (see **Table EV1** for model parameters).

We next evaluated the feedback loops, which are critical for the ‘closed loop’ architecture of the circuit, i.e., deactivation of Arf1 (mG*) by ArfGAP2/3 (mGAP) (**Figure 3H**). Two negative feedback loops activate ArfGAP2/3 (arrows 2 and 3 in **Figure 1A**). Arrow 2 represents GIV’s ability to bind and recruit ArfGAP2/3 to COPI vesicles and the Golgi membranes; failure to do so results in elevated levels of Arf1•GTP and stalled anterograde secretion in these cells (Lo et al., 2015). Arrow 3 represents GIV’s ability to activate Gi and release of ‘free’ Gβγ; GIV’s GEF function triggers this (Lo et al., 2015) and ‘free’ Gβγ is a co-factor for ArfGAP2/3. Both negative feedback loops depend on the forward reaction, arrow 1, which involves the recruitment of GIV (tGEF) (**Figure 1A**). Using the Arf1 activity after ligand stimulation as a readout, we next measured the activity of ArfGAP2/3 in control and GIV-depleted (i.e., shGIV) cells responding to EGF (**Figure 3I**). We found that, Arf1 activity peaked within 5 min after EGF stimulation and rapidly reduced thereafter by 15 min in control cells but remained sustained until 15 min in GIV-depleted cells (**Figure 3I, Figure EV2E-F**), suggesting that EGF activates both mGEFs and mGAPs of Arf1. While activation of Arf1 is brought on by mGEF(s) (the identity of which remains unknown) and achieves similar levels of activation regardless of the presence or absence of GIV, termination of Arf1 activity by mGAP (ArfGAP) requires GIV.

Finally, we used the model which was fitted to the experimental data in **Figure 2C** and **Figure 3G** to make predictions. We conducted two simulations: large decrease in the GIV level to simulate the Arf1 activation dynamics in shGIV cell (red line in **Figure 3J**), and delete either arrows 2 or 3 to simulate the Arf1 and Gi activation dynamics for the uncoupled GTPase switches. Based on the experimental results before (Lo et al., 2015), arrows 2 and 3 are modeled by an ‘AND gate’-like digital logical operation (Kime and Mano, 2003), i.e., a HIGH output (ArfGAP2/3 activity, and resultant termination of Arf1 signaling) results only if both the inputs to the AND gate (arrows 2 and 3) are HIGH. We also tested the ‘OR’ logic for the negative feedback (**Figure EV3**) and found the model predictions to be indistinguishable from AND gate. It is possible that one of these logical modes of operation is more efficient than the other under certain circumstances. For the first simulation, the simulated Arf1 activation dynamics (red line in **Figure 3J**) captured the sustained activation of Arf1 dynamics in shGIV cells, indicating the efficiency of the model. For the second simulation, the simulated Arf1 dynamics (green line in **Figure 3J**) is the same as that in shGIV cells, suggesting the equivalency of deleting GIV and uncoupling GTPase switch. The simulated Gi dynamics (green line in **Figure 3K**) is similar to (maybe even slightly higher than) that in control cells, which is consistent with the fact that the feedback loops have no effect on Gi. Thus, negative feedback within the ‘closed-loop control’ exerts significant effect on the mGTPase (Arf1) and little or no effect on the tGTPase (Gi).

### Coupled GTPases are predicted to enable high fidelity concordant response to EGF

To gain insights into how coupling impacts the information transduction, we compared the dose-response alignment (DoRA) performance between the coupled and uncoupled GTPase circuits. Typically, dose-response alignment (DoRA), referring to the close match of the receptor occupancy and the downstream molecules under different stimuli, is believed to improve information transduction, since the downstream molecules reflect the receptor occupancy faithfully. We regarded the mGEF as an alternative to the receptor because it serves as the first input to the coupled circuit via its ability to trigger the activation of the mGTPase switch. Therefore, a close match of dose-response curves of mGEF and mG* is equivalent to the linear relation between mGEF and mG*. Therefore, by using the model that has been fitted to the data in **Figure 2C** and **Figure 3G**, we simulated the steady-state value of mG* and mGEF over a wide range of stimulus, and then plotted the fractional activation of mG* for a given mGEF activity to observe the linearity. The misalignment in the case of a single switch is evident; a single Arf1 switch displays hyperresponsiveness, in that, max mG* is achieved even with minimal mGEF activity (**Figure 4A**). In the case of coupled switches, similar plots of fractional activation of mG* for a given mGEF activity show dose-response alignment with an unexpected linear relationship (**Figure 4B**). These results also hold in the presence of noise, such as noise in EGF stimulus and the intracellular noise [simulated within the concentrations of the different species (nodes) and the connections between them (arrows)] (see **Materials and Methods** and **Figure EV4**). These results suggest that coupled switches exhibit higher fidelity in information transduction than the uncoupled switches. Although unexpected for a GTPase switch, this finding is consistent with what is generally expected in a closed loop with negative feedback (Åström and Murray, 2021; Becskei and Serrano, 2000).

**Figure 4.**
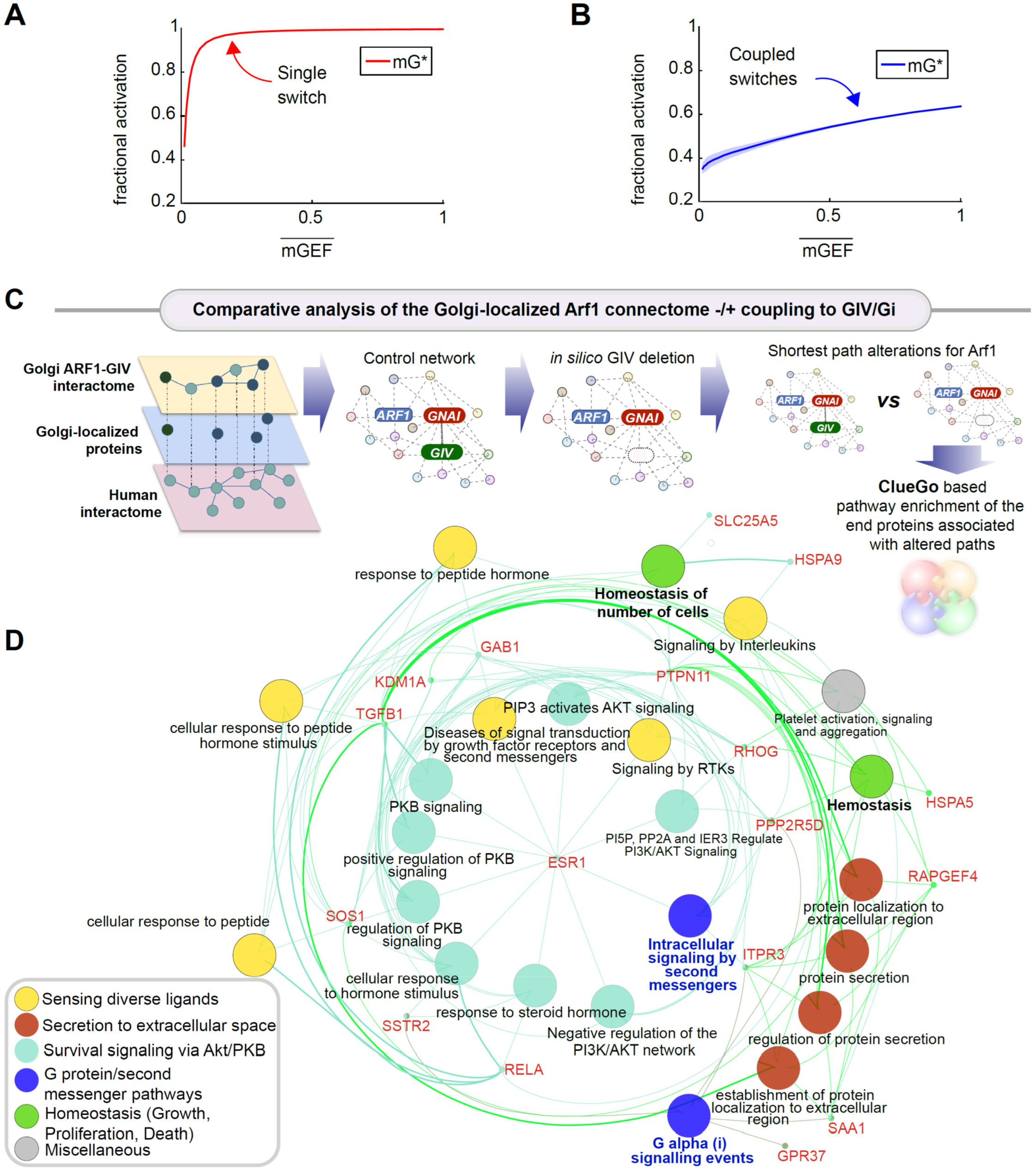
The predicted impacts of coupled switches on the degree of alignment of endomembrane responses (Arf1 activities; mG*) to the dose of extracellular stimulus, cellular processes, and fate. **A-B.** Fractional activations of mGEF versus active Arf1 (mG*) for the single switch (A; mG alone) and coupled switches (B; mG and tG). We perform stochastic simulations in the presence of noise in EGF (see **Materials and Methods** for details). The mean and the standard deviation (SD) of species are evaluated at steady states. 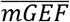 denotes the mean of mGEF; the shading shows the SD. The dimensionless EGF concentrations in the simulations are obtained through normalization, i.e., dividing the EGF concentration by 217.4 nM (= 50 nM/0.23). In all simulations, noise is introduced only in stimulus (i.e., EGF). **C-D.** A comparative analysis of the Golgi-localized Arf1 (mG) connectome with/without coupling to GIV (tGEF) and Gi (tGTPase). Workflow (C) shows how the list of Golgi-localized Arf1 and GIV interacting proteins (**Appendix Figure S1**; **Dataset EV1**) were used as ‘seeds’ to construct a PPI network from the STRING database to fetch the linking nodes to connect the seed proteins. The network was then perturbed by *in silico* deletion of GIV, followed by topological analysis of how such perturbation impacts the shortest paths associated with Arf1 to all other nodes in the network (see **Materials and Methods**). A network representation (**D**) using the ClueGo algorithm of the cellular processes associated with the end proteins that were most frequently encountered in the most impacted shortest paths associated with Arf1 (listed in **Appendix Figure S2E)**. The deleted or newly added shortest paths were only considered using the differential network approach (see **Materials and Methods**). The key on the lower left corner displays the color code of various overarching themes encountered in the network.

### Coupled GTPases are predicted to support secretion that is linked to autocrine signaling and survival

To understand the impact of uncoupling of the GTPase circuit on Arf1-dependent secretory functions of the Golgi, we carried out: (i) protein-protein interaction (PPI) network analysis and (ii) dynamical systems modeling.

To restrict the Arf1 interactome to the Golgi, we first extracted a Golgi-annotated subcellular localization network of high confidence GIV and Arf1 correlators, based on a proximity-dependent biotinylation map of a human cell (Go et al., 2021) (**Appendix Figure S1**). Next the list of Golgi-localized proteins was expanded by incorporating the GIV interactors from BioGRID (Oughtred et al., 2021) (**Appendix Figure S2A-B**). Arf1’s connectivity in the coupled network (in which Arf1•GIV•Gi interactions were intact) was compared against an uncoupled network created *in silico* by removal of GIV from the network (**Appendix Figure S2C**). Network analysis (see the workflow in **Figure 4C;** and as detailed in **Materials and Methods**) showed that Arf1’s connectivity with many proteins (‘nodes’) and pathways were altered in the uncoupled state (listed in **Appendix Figure S2D-G**). These altered pathways share three key themes: (i) “sensing” of diverse ligands/stimulus, e.g., growth factors, peptide and steroid hormones, cytokines (yellow nodes in **Figure 4D**), (ii) “secreting” proteins to the extracellular space (red nodes in **Figure 4D**) and (iii) “survival” signaling via the PI3K-Akt pathways (teal nodes in **Figure 4D**). As anticipated in the absence of GIV, Gi and second messenger signaling (blue nodes in **Figure 4D**), cellular homeostasis and cell number (green nodes in **Figure 4D**) were predicted to be impacted. These findings suggest that removing GIV may impact secretion that is critical for auto/paracrine sensing/signaling, which maintains cell number via balanced proliferation and/or death.

We next used systems modeling approaches to interrogate how coupled (closed loop control) *vs*. uncoupled (open loop) GTPase systems at the Golgi impact cargo secretion and cell number upon sensing growth factor stimulus (**Figure 5A**). However, unlike m/tG* activity assays (which happen in sec to min), cell secretion may begin within min but is measured in hours, and their impact on cell number requires a longer time scale (several hours and even, days). The secretion function was predicted to show an ultrasensitive response (n_Hill_=1.86) as a function of the stimulus (i.e., a given dose of EGF) when the two GTPase switches are coupled; it was predicted to be reduced in the absence of coupling (**Figure 5B-D**). Though the secretion response is a constant in the absence of the coupling, it sets the baseline for EGF-to-Arf coupling and thus supplies a platform for the comparison with the coupled GTPases system. Intriguingly, secretion in the coupled state shows different response for most ranges of Gi activity (tG*; **Figure 5C**), indicating a faithful information transduction between Gi and secretion. Furthermore, the cell number is higher for the coupled versus the uncoupled cells (**Figure 5E** and **Appendix Figure S3**). To specifically analyze the impact of secrete-and-sense autocrine autonomy, we carried out the simulations under restrictive growth conditions. These simulations under restrictive growth conditions revealed that cells with coupled switches display a higher cell number compared to the cells with uncoupled switches only when the secrete-and-sense loop is highly efficient; this advantage is lost if the loop is abolished (**Figure 5F**). That coupling of GTPases is required for maintaining cell numbers was reproduced using EGF as stimulus (**Figure 5G**), providing continuity with prior model-derived predictions. We also confirmed that the system and the conclusions are not only robust to biological noise (**Figure 5B, C, G**), but also robust to the variations in the kinetic parameter (**Appendix Figure S4**).

**Figure 5.**
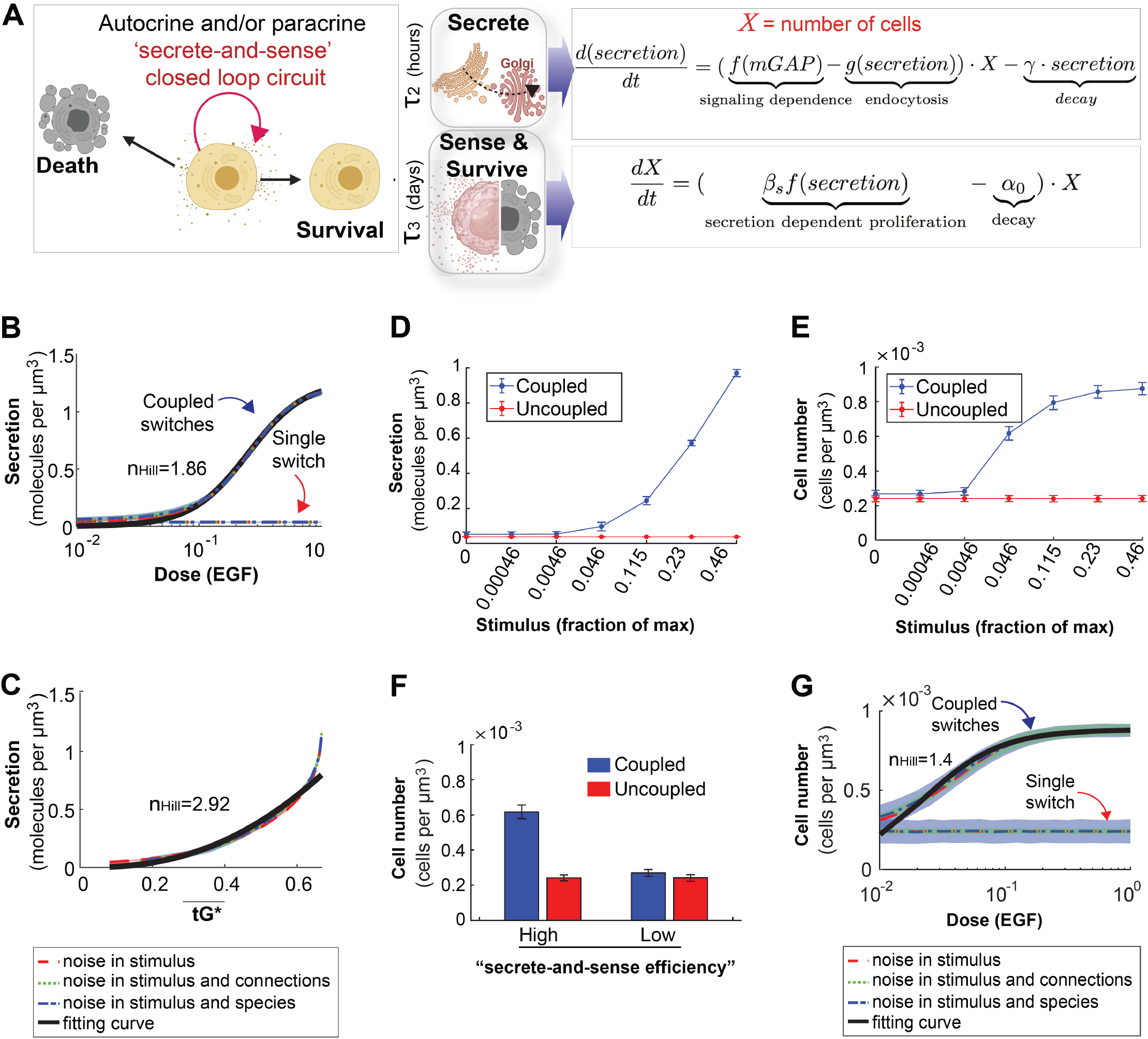
Coupled GTPases are predicted to support secrete-and-sense autonomy and for the maintenance of cell number. **A.** Schematic of the key features of the auto/paracrine loop that we hypothesize is regulated by the coupled GTPase circuit (left) and the corresponding phenomenological models to capture these key effects (right). **B-C.** Model prediction for secretion as a function of stimulus in cells with coupled and uncoupled GTPases. Noise is introduced into the system in a similar way as described in **Figure EV 4D-G**. r2>0.99 in B; r2>0.94 in C. **D-E.** The secretion (D) or the cell number (E) as a function of stimulus in coupled and uncoupled switches. The stimulus =0, 0.00046, 0.0046, 0.046, 0.115, 0.23 and 0.46 correspond to varying doses of EGF in simulations, ranging from 0, 0.1 nM, 1 nM, 10 nM, 25 nM, 50 nM, and 100 nM, respectively. The error bar denotes the S.D when noise is in the stimulus and connections. **F.** The bar plot depicts cell numbers achieved by cells with either coupled or uncoupled switches, at different levels of stimulus. The first two bars represent the cell number when stimulus =0.046, and the last two bars are for stimulus =0. Noise is introduced as in B-C. **G.** Relation between cell number and EGF in the presence of noise, which was introduced in a similar way as described in **Figure EV4 D-G**. r^2^>0.95.

### GTPase coupling by GIV is required for time and dose-dependent secretion of diverse cargo proteins

We next sought to experimentally validate the predicted impact of uncoupling on cell secretion by studying the time-dependent secretion of a few well-established transmembrane and soluble cargo proteins. We began with the transmembrane cargo, vesicular stomatitis virus G protein (VSVG) using the well-characterized GFP-tagged VSVG-tsO45 mutant (Gallione and Rose, 1985). This mutant VSVG is retained in the ER at 40°C, accumulates in Golgi stacks at 20°C temperature block, from where it escapes to the PM at permissive 32°C (**Figure EV5A**). Considerable VSVG accumulated in the Golgi region in both control and GIV-depleted cells under serum-starved conditions at 20°C. EGF or serum stimulation was permissive to transport of the VSV-G protein to the PM in control cells at 32°C, but such transport was significantly diminished in GIV-depleted cells (**Figure EV5B-C**). Similar results were observed also in the case of EGF-stimulated secretion of three separate soluble cargo proteins, MMP2, MM9 (**Figure EV5D-F**) and Collagen (**Figure EV5G-H**); these cargo proteins were chosen because of GIV’s published role in ECM degradation during cancer metastasis (Rahman-Zaman et al., 2018) and tissue fibrosis (Lopez-Sanchez et al., 2014). These findings show that the secretion of diverse proteins in response to growth factors is blunted in GIV-depleted cells with uncoupled GTPases (**Figure 6A**).

**Figure 6.**
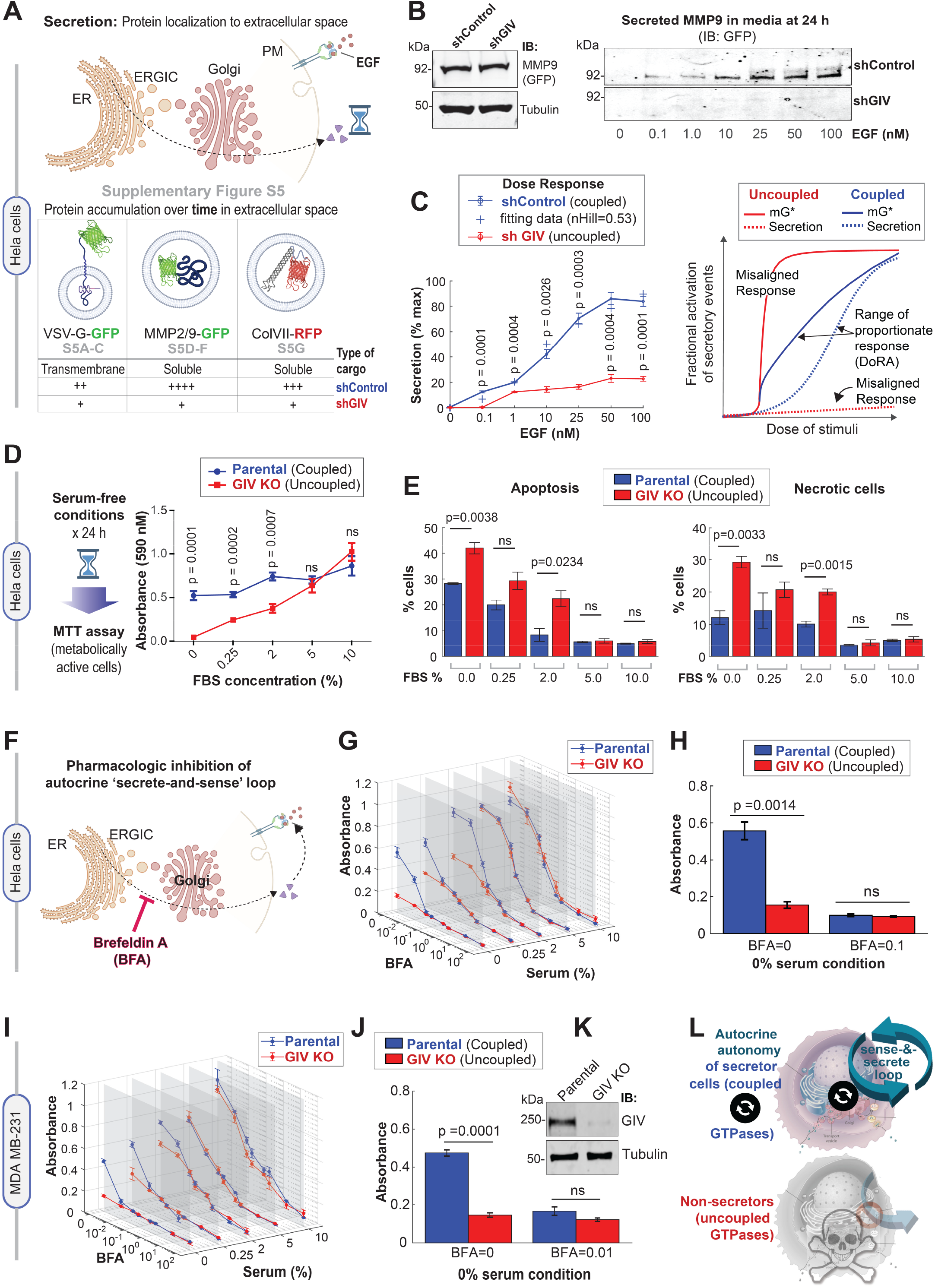
Coupling of GTPases by GIV is required for growth factor-independent cell survival that relies upon autocrine secretion. **A.** Schematic summarizes the findings showcased in **Figure EV5**, which investigate the secretion of diverse cargo proteins [temperature-sensitive (ts) VSV-G, MMP2/9 and ColVII], as determined by their accumulation in extracellular space over time after stimulus (EGF or serum). For each cargo tested, compared to cells with GIV (shControl), ligand-stimulated secretion was impaired in cells without GIV (shGIV). **B**. Immunoblots showing intracellular (left) and secreted (in the media; right) GFP-MMP9 at 24 h after stimulation with varying doses of EGF. Tubulin, used as loading control, confirm the presence of a similar number of plated cells in the assay. **C**. *Left:* Graph displays experimentally determined secretion of GFP-MMP9 in response to varying doses of EGF in control (shControl) and GIV-depleted (shGIV) HeLa cells (as in B), and quantified by band densitometry. Results are expressed as mean ± S.E.M; n = 3. p values were determined by two-sided unpaired t-test. *Right*. Schematic diagram of dose responses (mG* and secretion) for the single switch and coupled switches. Coupled switches stretch the range of proportionate responses. Single mG switch results in misaligned responses. DoRA, dose response alignment. **D.** Left: Schematic summarizing the colorimetric assay used here to determine the number of metabolically viable cells. Right: The graph displays formazan absorbance expressed as a measure of cell viability from the HeLa cells (Y axis) cultured at varying conc. of serum (X axis). Results are expressed as mean ± S.E.M; n = 3. p values were determined by two-sided unpaired t-test. **E.** Bar graphs display the % apoptotic (*left*) or necrotic (*right*) control (parental) and GIV-depleted (GIV KO) HeLa cells after 24 h growth in varying concentrations of serum, as assessed by annexin V staining and flow cytometry. See also **Appendix Figure S5A-C** for dot plots and early and late apoptotic fractions. Results are expressed as mean ± S.E.M; n = 3. p values were determined by two-sided unpaired t-test. **F.** Schematic showing the rationale for and mechanism of action of fungal toxin, BFA, for interrupting the secrete-and-sense autocrine loop in cells. **G-H.** Control (parental) and GIV-depleted (GIV KO) HeLa cells grown in different concentrations of serum (FBS%) were treated or not with varying concentrations of BFA (μM) as indicated. Line graphs in 3D (G) depict the formazan absorbance expressed as a measure of cell viability from the HeLa cells in various conditions tested. Bar graphs (H) depict the cell number in serum-free growth conditions that are supported exclusively by autocrine secrete-and-sense loop (without BFA; BFA = 0.0 μM) or when such loop is interrupted (BFA = 0.1 μM). Results are expressed as mean ± S.E.M; n = 3. **I-K**. Control (parental) and GIV-depleted (GIV KO) MDA MB-231 cells grown in different concentrations of serum (FBS%) were treated or not with varying concentrations of BFA (μM) as in G-H. Line graphs in 3D (I) depict the formazan absorbance expressed as a measure of cell viability from the MDA MB-231 cells in various conditions tested. Bar graphs (J) depict the viability of the MDA MB-231 cells in serum-free growth conditions that are supported exclusively by autocrine secrete-and- sense loop (without BFA; BFA = 0.0 μM) or when such loop is interrupted (BFA = 0.1 μM). Immunoblots (K) of equal aliquots of whole cell lysates confirm the depletion of GIV compared to tubulin (loading control). See also **Appendix Figure S5D-H** for dot plots and early and late apoptotic fractions. Results are expressed as mean ± S.E.M; n = 3. **L**. Summary of conclusions of this work. *Top*: Coupling of GTPases within the secretory pathway enables dose-response alignment of secretion to stimulus, which appears to be essential for ‘secrete-and-sense’ autocrine autonomy in cancer cells. *Bottom:* Uncoupling of the GTPases within the secretory pathway disrupts such autonomy and leads to cell death.

We next asked if the DoRA predicted earlier in the case of Arf1 activity (**Figure 4B**) translates into a similar alignment in the case of cell secretion. We analyzed the efficiency of secretion of one of the 3 cargo proteins, MMP9, from control and GIV-depleted cells responding to a range of EGF concentrations for 24 h (**Figure 6B**). Quantitative immunoblotting confirmed that dose-dependent secretion was observed in the case of control cells (coupled GTPases) but not in GIV-depleted cells (uncoupled GTPases) (**Figure 6B; 6C** *left*). We conclude that DoRA of Arf1 activity indeed translates into DoRA of cell secretion in cells with coupled GTPases; by contrast, a misaligned Arf1 activity (hyperresponsive; **Figure 4A, 6C**, right) translates into misaligned secretion (hyporesponsive; **Figure 6C**, right) in cells with uncoupled GTPases.

### GTPase coupling by GIV is required for cell survival that relies upon autocrine secretion

We next assessed by MTT assays the total number of metabolically active cells that develop self-sufficiency in growth factor signaling, i.e., survive in growth factor-free conditions (0% serum) (**Figure 6D**, *left*). The number of cells in serum-free or low-serum conditions was significantly higher in the presence of GIV (parental HeLa cells; coupled) than in the absence of GIV (GIV-KO cells; uncoupled) (**Figure 6D**); this survival gap closed at higher serum concentrations (see 10% FBS, **Figure 6D**). Reduced cell number in GIV-KO cells in the low/no serum conditions was associated with a concomitant increase in cell death via apoptosis and necrosis (**Figure 6E, Appendix Figure S5A-C**). We then sought to validate the results of the simulations in growth-restrictive conditions which showed that interrupting the coupled GTPase circuit at the Golgi will reduce cell numbers (**Figure 5F**). We analyzed the number of metabolically active cells with (coupled) or without (uncoupled) GIV across a range of serum conditions and varying concentrations of the mycotoxin Brefeldin A (BFA), a well-known tool to inhibit secretion via its ability to inhibit Arf1 activation (Prieto-Dominguez et al., 2019) (**Figure 6F**). We made three observations: (i) cells with coupled circuits have a significant survival advantage in serum-restricted conditions (see 0 - 2.0% FBS; **Figure 6G**); (ii) that advantage depends on sensing what the cells secrete, because blocking secretion with BFA also eliminates such advantage (**Figure 6H**); (iii) survival in the presence of serum (5-10%) is similar for both “coupled” and “uncoupled” cells, implying non-secreting cells with uncoupled circuits can survive if they can “sense” stimuli that they did not generate (e.g., serum ~5-10% range; **Figure 6G**). Cells with coupled circuits have a significant survival advantage in serum-restricted conditions (see 0 - 2.0% FBS; **Figure 6G**); (ii) that advantage depends on sensing what the cells secrete, because blocking secretion with BFA also eliminates such advantage (**Figure 6H**); (iii) survival in the presence of serum (5-10%) is similar for both “coupled” and “uncoupled” cells, implying non-secreting cells with uncoupled circuits can survive if they can “sense” stimulus that they did not generate (e.g., serum ~5-10% range; **Figure 6G**).

To avoid overreliance on a single cell line (i.e., HeLa), we generated a second model, GIV-depleted MDA MB-231 cells (by CRISPR, see **Materials and Methods**) and sought to reproduce key findings (**Figure 6I-K**). As in the case of HeLa cells, the survival advantage of MDA MB-231 cells with coupled circuit (with GIV, Parental cells) over those with uncoupled circuit (GIV KO) was observed exclusively in low/no serum conditions (see 0 - 2.0% FBS; **Figure 6I**) and blocking secretion with BFA eliminates such advantage (**Figure 6J**). Reduced cell survival in cells without GIV (uncoupled state) was associated with higher early and later apoptosis and necrosis (**Appendix Figure S5D-H**).

These findings show that the coupled GTPase circuit is required for cell survival that is supported exclusively by autocrine secretion (i.e., independent of external growth factors), and by that token, essential for a functional autocrine ‘secrete-and-sense’ loop (**Figure 6L,** *top*). Interrupting the coupled GTPase circuit at the Golgi appears to disrupt the ‘secrete and sense’ loop and abrogate cell survival that is supported by such secretion (**Figure 6L,** *bottom*). Because ‘secrete-and-sense’ loop is a key feature of cellular autonomy (Maire and Youk, 2015; Youk and Lim, 2014), taken together our findings show that the coupled GTPase circuit in the cell’s secretory pathway may be critical for autocrine autonomy.

### GTPase coupling supports self-sufficiency in growth factor signaling

To discern the nature of the pathway/processes whose autocrine autonomy is supported by the coupled GTPases, we analyzed HeLa and MDA MB-231 cells with coupled (WT) or uncoupled (GIV KO) circuits by tandem mass tag (TMT) proteomics. The studies were carried out in serum-free/restricted conditions (**Figure 7A**) to maximally enrich the proteome that supports auto-/paracrine secretion-coupled sensing. To our surprise, the majority (76%; 1437 proteins, includes EGFR; see complete list in **Dataset EV2**) of the differentially upregulated proteins (DEPs) in the two WT cell lines overlapped (despite the vast differences between HeLa and MDA MB-231 cell lines in origin, genetics, and nearly every other possible way). This suggests that the presence or absence of GTPase coupling via GIV may impact both cells similarly. The interactions between the DEPs were fetched from the STRING database to build a PPI network, which showed major coat proteins (AP1, AP2, COP, and CAV), monomeric GTPases (Arfs, Rabs, Rho, CDC42 and Rac1) and trimeric GTPases (GNAI) (**Figure 7B**). A degree of connectivity analysis revealed that EGFR and the Arfs are some of the most highly connected nodes in the interactome (**Figure 7C**). A reactome pathway enrichment analysis confirmed that the most highly connected proteins primarily engage in a variety of growth factor signaling pathways (**Figure 7D**).

**Figure 7.**
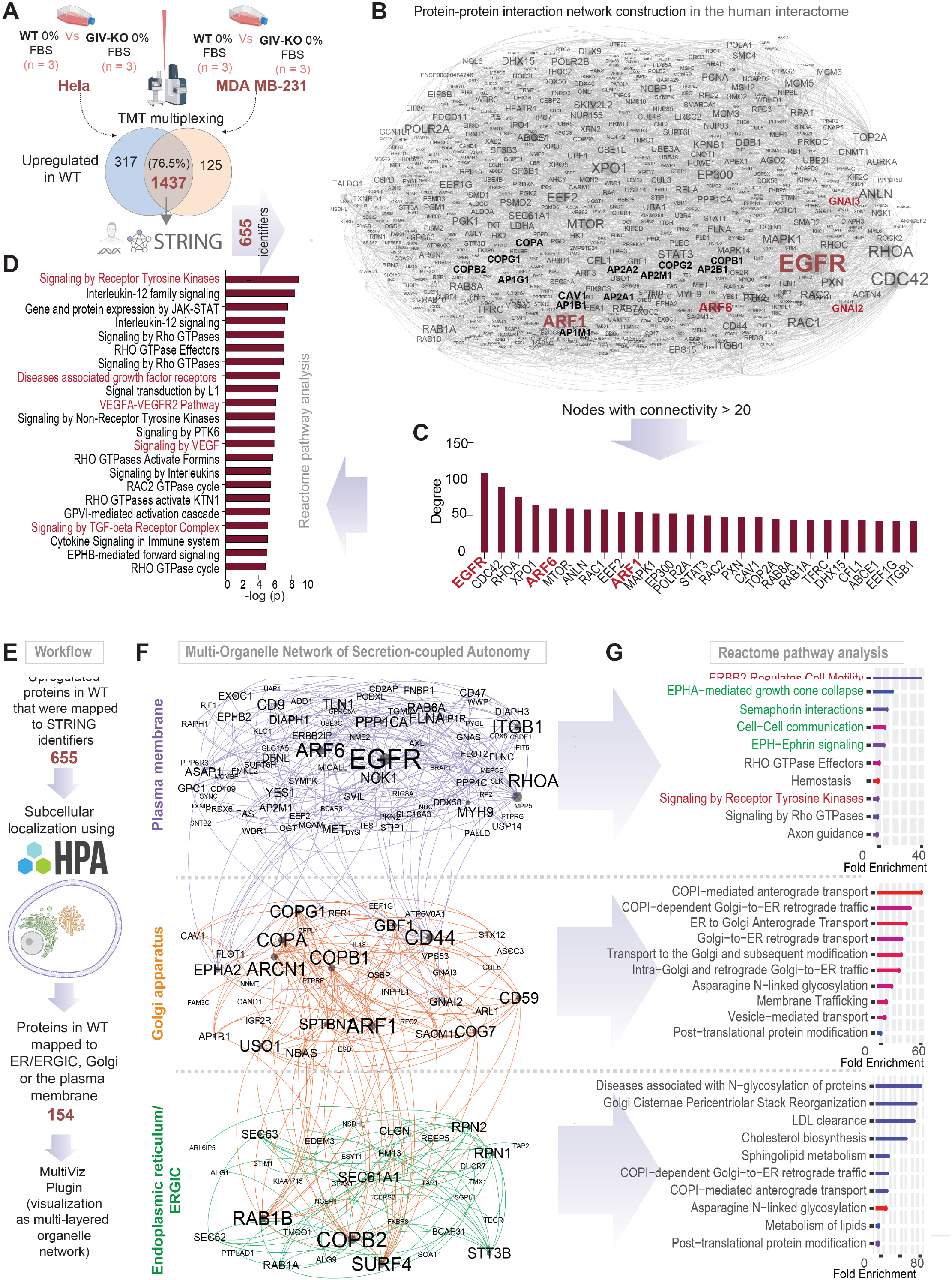
Differential proteomics of autonomy enabled versus disabled MDA MB-231 and Hela cells. **A.** Workflow for comparative proteomics on autonomy enabled vs. disabled cells by tandem mass tag (TMT) multiplex technique followed by mapping of upregulated proteins in WT cells using the STRING database (see **Materials and Methods**). **B.** A protein-protein interaction (PPI) network shows the interactions between upregulated mapped proteins in WT cells. Node and font sizes correlate positively with the degrees of connectivity. **C.** Bar plot shows the degree distribution of highly connected (degree >20) nodes in the PPI network in B. **D.** Reactome pathway analysis of the pathways enriched in the most connected proteins in C. Red = pathways associated with growth factor signaling. **E.** Workflow for the construction of a multi-organelle network of autonomy enabled cells using subcellular localization of upregulated proteins in the WT cells. **F.** Visualization of a multi-organelle network of proteins that partake in secretion-coupled autonomy across 3 compartments, the plasma membrane, the Golgi and the ER/ERGIC. **G.** Reactome pathway analyses of the pathways enriched within the three organelles in F. Red pathways associated with RTK/EGFR signaling and Green pathways associated with multi-cellular cell-cell communication in the plasma membrane.

Because protein functions are determined by subcellular localization, we sought to map the DEPs that are upregulated in the WT cells based on their subcellular localization. To this end, we used a Human Cell Atlas-supported explorer platform that was generated using the large collection of confocal microscopy images showing the subcellular localization patterns of human proteins to include in analysis three organelles, the Golgi, ER/ERGIC and the PM (**Figure 7E**; see **Materials and Methods**). Visualization of the DEP derived PPI network as multi-layered networks that are comprised of intra- and inter-organelle interactions (**Figure 7F**) revealed greater insights. As expected, reactome pathway analysis of the ER/ERGIC and the Golgi interactomes showed an enrichment of protein processing and secretory processes, respectively, and the PM-localized interactome showed an enrichment of growth factor signaling (**Figure 7G**). The PM-localized interactome also showed an enrichment of cell-cell contact and contact-dependent signaling pathways (such as Semaphorins and the Eph-ephrin system; green; **Figure 7G**), which enable cell-cell coordination in multicellular eukaryotes. These findings indicate that the coupled GTPase system supports a network of proteins that primarily enable secretion-coupled growth factor sensing and thereby, growth signaling autonomy.

## DISCUSSION

The major discovery we report here is the creation of an experimentally constrained multi-timescale model for cell survival that relies on growth factor-responsive cell secretion. One major consequence of such a phenomenon is autonomous growth/survival in the absence of external growth factors. We formally define the molecular basis for such autonomy and demonstrate the consequences when it is manipulated/perturbed. The insights and models derived from this study are expected to inform and impact at least three fields, i.e., of signal transduction, cell secretion, and cancer cell biology in the following ways.

In the field of signal transduction, emergent properties of ectomembrane signaling circuits at the PM have been identified using systems biology; however, none thus far have coupled the events at the ectomembrane to the events in the cell’s interior, i.e., the endomembrane of organelles. Our study experimentally validated a Golgi-localized natural coupling between the two GTPase switches with exquisite feedback control that enables linear activation of Arf1 in response to EGF, which in turn enables the Golgi to mount a response (protein secretion) that is proportionate to the stimulus (sensed at the PM) and robust to noise. The model reveals two notable features: First we show that the closed loop control system generated DoRA, enabling a linear increase in Arf1/mG* activation and protein secretion. Such DoRA has been described in several major receptor-initiated signaling cascades at the PM (from the pheromone response system in yeast to the Wnt→βCatenin, TGFβ→SMAD2/3 and EGFR→MAPK cascades in mammals) (Andrews et al., 2016), but never in endomembrane GTPases. Because a linear DoRA maximally preserves any information during its propagation (Andrews et al., 2018), we conclude that one of the major discernible consequences of the closed loop coupling of two GTPases is its ability to faithfully transmit information from the PM to the Golgi for the latter to mount a concordant secretory response. Second, although the first switch, i.e., Arf1/mG* showed a linear response, the subsequent steps (switch #2 and the step of membrane mechanics leading to secretion) become progressively ultrasensitive. The net result of this is that the closed loop feedback control allows for a tighter alignment of secretion with respect to EGF by ‘stretching’ out the dose-response curve across series of switches to propagate the signal from the extracellular space to the interior of the cell. Because the stability behavior of a mathematically simpler version of this closed loop system of coupled GTPases showed that coupling afforded a wide range of steady states (Stolerman et al., 2021), it is tempting to speculate that the coupled system allows flexibility in responses over a wide range of stimulus. In fact, follow-up work has now revealed how ranges of activity of the mGTPase Arf1, reaction kinetics, the negative feedback loop (mGAP), and the cascade length affect DoRA (Qiao et al., 2022).

When it comes to the field of protein secretion, the cell’s secretory pathway was originally believed to be a constitutive function that is regulated by ‘housekeeping’ genes/proteins that maintain the integrity of the local (membrane or lumenal) environment (Arvan et al., 2002). The earliest evidence that secretion is regulated by exogenous growth factors emerged in 2008 when the phosphoinositide phosphatase SAC1 was implicated as a ‘brake’ in anterograde Golgi secretion that is released by growth factors (Blagoveshchenskaya et al., 2008). Despite these insights, what remained unknown was how the secretory system (or any intracellular organelle/system) responds proportionately to the external cues. The functional consequences of an endomembrane coupled GTPase system we dissected here fill that knowledge gap. We show that coupling of m/tGTPases with closed loop control within cells is critical to set up feedback controls in yet another scale, i.e., cell secretion and cell fate (i.e., survival *vs*. death).

Finally, when it comes to the field of cancer cell biology, it is well accepted that self-sufficiency in growth signaling is a hallmark of all cancer cells (Hanahan and Weinberg, 2000); we show here how cells achieve such self-sufficiency for the prototypical growth factor system, i.e., EGF/EGFR. Existing theories linking genetic circuits to cellular autonomy, although quantifiable and tunable (Doğaner et al., 2016; Kamino et al., 2017; Maire and Youk, 2015; Tang et al., 2021; Youk and Lim, 2014), don’t apply to the multicellular eukaryotes. In dissecting the behavior of the coupled GTPase system, and revealing the consequences of its disruption, both *in silico* and in two different cancer cells, we fill that knowledge gap. Second, intratumoral cellular heterogeneity is known to give rise to an ecosystem of clonal interactions (Basanta and Anderson, 2013; Tabassum and Polyak, 2015) that can drive tumor growth, therapeutic resistance, and progression (Basanta and Anderson, 2017; Li and Thirumalai, 2019; Maley et al., 2017; Merlo et al., 2006). Therefore, it is possible that a few autonomous secretor clones with an intact secrete-and-sense loop could be sufficient to support the survival of neighboring non-secretor clones. If so, uncoupling the GTPases and disrupting the secrete-and-sense autonomy could serve as an impactful therapeutic strategy. Finally, the evolutionary significance of our findings is noteworthy. For example, the linker between the GTPases, i.e., GIV, evolved later in multicellular organisms such as worms (Nechipurenko et al., 2016) and flies (Ha et al., 2015; Houssin et al., 2015; Puseenam et al., 2009; Yamaguchi et al., 2010). GIV’s HOOK module (binds mGTPase) evolved in worms and flies (Ha et al., 2015; Houssin et al., 2015; Puseenam et al., 2009; Yamaguchi et al., 2010); its GEM domain (a short motif that binds and modulates tGTPases) evolved later in fish (DiGiacomo et al., 2018) and remains to date. Thus, the coupled GTPase circuit likely evolved in higher eukaryotes, and as suggested by our multi-organelle proteomic analyses, is geared to support autonomy in multicellular organisms. This is consistent with the fact that evolution appears to favor efficient signaling circuits that can accomplish many different tasks (Milo et al., 2002; Shen-Orr et al., 2002). Because GIV is overexpressed in the most aggressive tumor cells, it is likely that the GTPase coupled circuit is more frequently assembled in those cells. If so, the circuit may represent an evolutionary masterpiece of multiscale feedback control to achieve autonomy, adaptability, and flexibility. Follow-up work has now shed light on the importance of this phenomenon in the orchestration of self-sustained EGFR/ErbB signaling in tumor cells (Sinha et al., 2022). Such autonomy in growth signaling appears to be critical for the maintenance of high metastatic potential and epithelial mesenchymal plasticity during the blood-borne dissemination of human breast cancer.

## LIMITATIONS OF THE STUDY

The multi-timescale model we built, ignores the spatial aspects of the various feedback control loops. Because the spatial organization of signaling motifs will influence their temporal behaviors, we anticipate the need for further refinement of the current model. By depleting GIV, we disconnect the GTPases and dismantle the entire circuit; selective disruption of various connections within the Golgi-localized circuit is not possible currently due to the lack the experimental tools (e.g., specific point mutants of GIV, GEF or GAPs or perturbagens such as small molecule or peptides. Although we studied four different cargo proteins (VSV-G, MMP2/9 and Col-VII) and two types of stimuli (EGF and serum), a more comprehensive assessment of the cell’s secretome is expected to reveal how the intracellular GTPase circuit controls the composition of the extracellular space. We chose to model to test the experimentally determined key components by design, but there may be missing components which enable other emergent properties (such as, advantages of AND vs OR gate mechanisms in the feedback loops); future work is expected to build upon this framework to fill these knowledge gaps. Conducting experiments across the full range of stimuli to assess ‘proportionality/linearity’ of response was possible in some instance (e.g., cell survival) but not possible in others (e.g., FRET, Arf1 activity, etc.) due to technical limitations of the assays and/or detection thresholds. Finally, our mathematical model ignores the effect of the physical location and heterogeneity of cells. To explore such homogeneous and heterogeneous cell population (Gerlee and Anderson, 2008; Poleszczuk et al., 2015; Sottoriva et al., 2010) future studies will need to include agentbased models (Chao Dennis et al., 2008; Norton and Popel, 2014; Wang et al., 2007), in which each cell is regarded as an individual agent that ‘senses’ the environment and ‘decides/acts’ in response.

## Abbreviations

GIV: Gα-interacting Vesicle-associated protein
GEM: guanine nucleotide exchange modulator
DoRA: dose-response alignment
Arf1: ADP-ribosylation factor-1
EGF: epidermal growth factor
ER: endoplasmic reticulum
ERGIC: ER-Golgi Intermediate Compartment
CLC: closed loop control
GGA: Golgi-localized, γ-ear-containing, Arf -binding protein

## AUTHOR CONTRIBUTIONS

L.Q carried out and analyzed all modeling studies in this work. S.S carried out TMT proteomics studies, proteomics analysis, and visualization of data as multi-layered network. A.A.A., I-C.L, and C.R designed, carried out and analyzed most of the experiments and assembled the figures. T.N extracted the Golgi-specific proteome from the human cell map and performed the GO analyses; SS performed the protein-protein network analyses. V.G. performed VSVG transport assays; N.A and I-C.L performed the MMP and collagen secretion assays. K.M. and I.L-S carried out and analyzed the FRET and Gαi:GTP assays, respectively. L.Q, AAA, I-C.L and V.G. helped write methods and edit the manuscript. P.G. and M.G.F conceptualized, designed, supervised and analyzed the experiments. P.R, with input from P.G, conceptualized, designed, supervised and analyzed the modeling studies. P.G, P.R, and L.Q wrote the manuscript.

## ACKNOWLEDGEMENTS

We thank Linda Joosen for technical assistance with FRET assays. This work was supported by the following National Institute of Health Grants: CA238042, AI141630 and CA160911 (to P.G.), GM132106 (to P.R), CA100768 (to M.G.F and P.G). Other sources of support include the Wu Tsai Human Performance Alliance and the Joe and Clara Tsai Foundation. P.R was also supported by the Air Force Office of Scientific Research (AFOSR) Multidisciplinary University Research Initiative (MURI) Grant FA9550-18-1-0051. I-C. L. was supported in part by a Fellowship (NSC 100-2917-1-564-032) from the National Science Council of Taiwan, K.M. by Susan G. Komen award (# PDF14298952) and I.L-S by the American Heart Association (AHA #14POST20050025).

A.A.A was supported, in part, by an NIH-funded Cancer Therapeutics Training Program (CT2, T32 CA121938).

---

**Table EV1:**
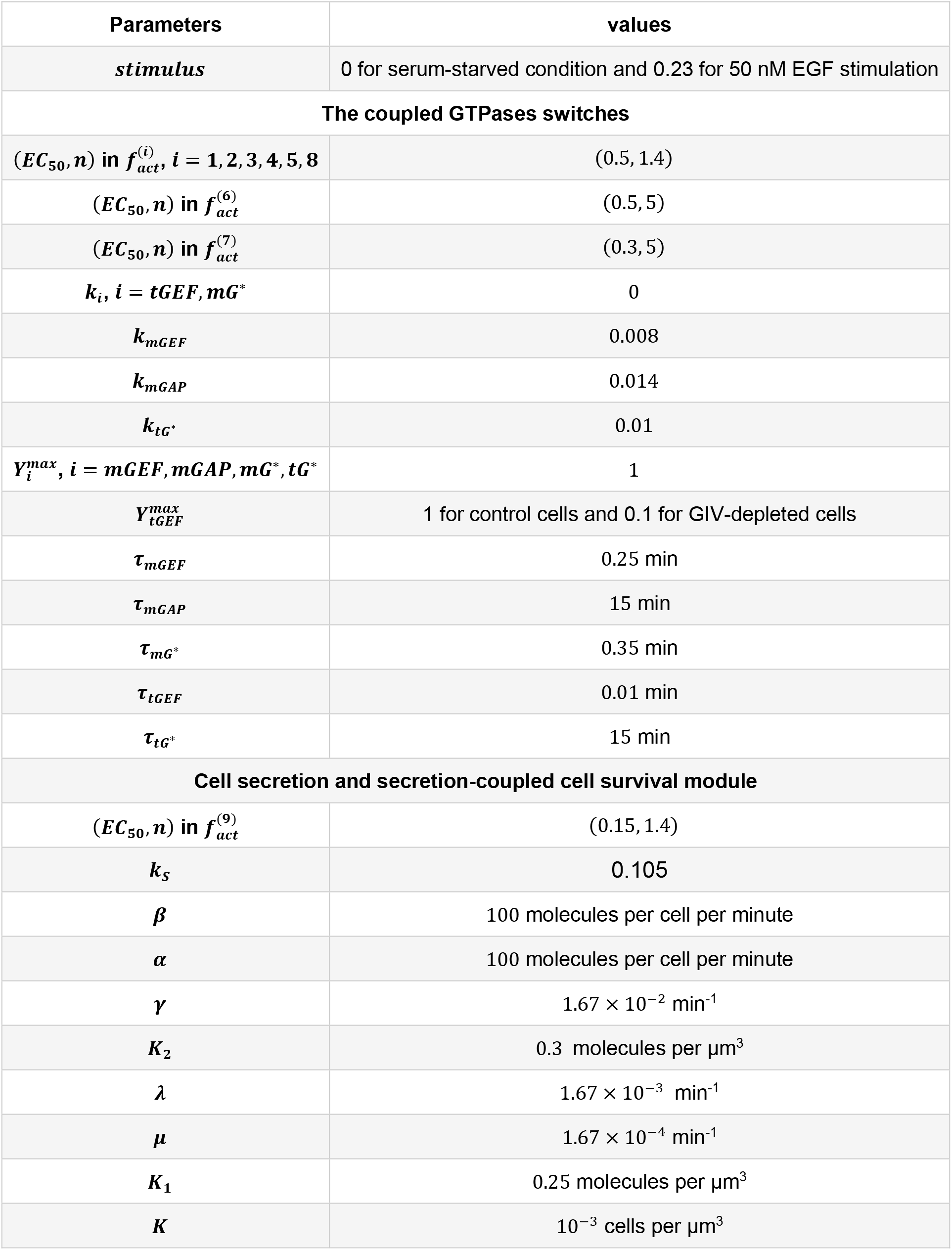
Parameter values used in the deterministic model

## EXPANDED VIEW (EV) FIGURES

**Figure EV1.**
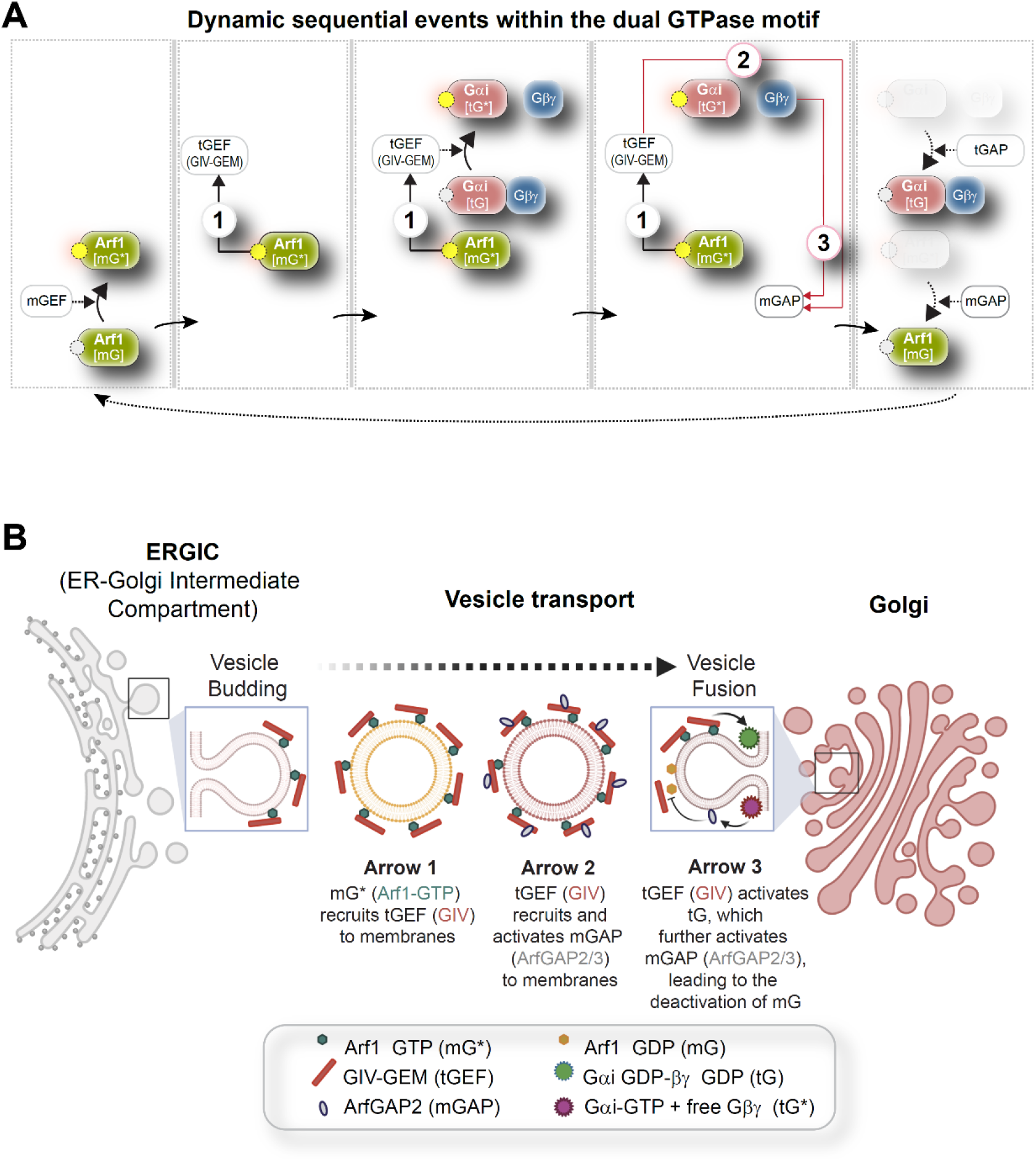
An endomembrane network motif of two species of GTPases regulates membrane trafficking through the secretory pathway, regulates Golgi functions. **A.** Dynamics within the endomembrane GTPase system. Left to right panels display the deconstructed Arrows denote key molecular events/chemical reaction cascades within this system, in which, the GIV-GEM links monomeric (m) and trimeric (t) GTPase systems and enable the conversion of extracellular stimulus (ligand; left) into membrane trafficking events (e.g., vesicle uncoating/budding/fusion; right). The forward and feedback reactions (arrows) are numbers 1-3. See **Movie EV1** for a gif of the circuit. **B.** Schematic summarizing the findings reported by Lo. I., et al., (Lo et al., 2015) delineating how arrows 1-3 within the endomembrane GTPase system regulates finiteness of Arf1 signaling for efficient secretion.

**Figure EV2.**
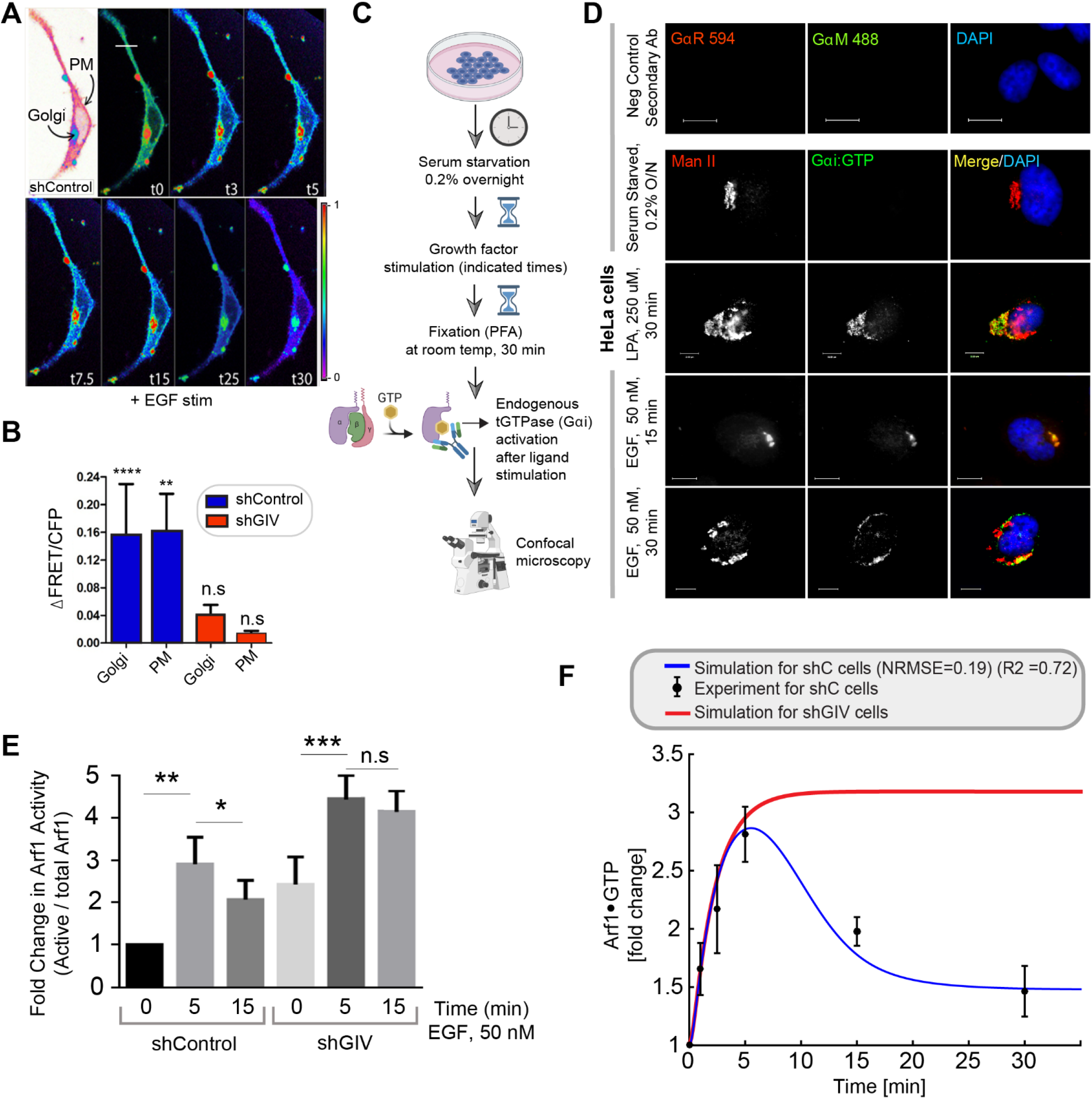
GEV-GEM is required for Gi activation at the Golgi and for maintaining the finiteness of Arf1 signaling upon EGF stimulation. **A.** FRET based studies were carried out in sh Control cells as in **Figure 3B-C**. Briefly, HeLa cells were co-transfected with Gαi1-YFP, Gβ1-CFP and Gγ2 (untagged) and live cells were analyzed by FRET imaging at steady-state, after being serum starved in 0.2% FBS overnight and then after stimulation with 50 nM EGF. Representative freeze-frame FRET images are shown. FRET image panels display intensities of acceptor emission due to efficient energy transfer in each pixel. FRET scale is shown in inset. Golgi and PM regions of interest are indicated with arrows. **B.** Bar graphs in I display the change in FRET at t5 min at the Golgi and the PM regions of 3-5 cells, from 4 independent experiments. Scale bar = 7.5 μm. Line graphs in J represent the dynamic change in FRET in the Golgi regions in sh Control vs shGIV cells. Results are displayed as ± S.E.M. **C.** Schematic showing how a conformation-specific anti-Gαi•;GTP antibody detects GTP-bound active Gαi in situ. **D.** HeLa cells starved with 0.2% FBS overnight or stimulated subsequently with 50 nM EGF or 250 μM LPA were fixed and stained for active Gαi (green; anti-Gαi:GTP mAb) and Man II (red) and analyzed by confocal microscopy. Activation of Gαi was detected exclusively after LPA/EGF stimulation. When detected, active Gαi colocalizes with Man II (yellow pixels in merge panel). Negative control (secondary antibody) staining was carried out on cells stimulated with EGF, 15 min. Scale bar = 7.5 μm. **E.** Control (sh Control) and GIV-depleted (shGIV) HeLa cells that were stimulated with EGF for the indicated time points prior to lysis were assessed for Arf1 activity. Immunoblots are shown in **Figure 3I**. Bar graphs display the fold change in Arf1 activity normalized to t0 min that was observed in control (shControl) and GIV-depleted (shGIV) cells. Results are expressed as mean ± S.E.M; n = 3 replicates; *p* values were determined using Mann-Whitney t-test compared to t0: *, <0.05; **, <0.01; ***, <0.001. Immunoblots are representative of findings from at least 3 independent repeats. **F.** Line graph in red displays model-derived simulation of Arf1 activation dynamics (mG*) in cells without tGEF (shGIV). As reference, results of model-derived simulation fit to experimental data in control cells are displayed in blue.

**Figure EV3.**
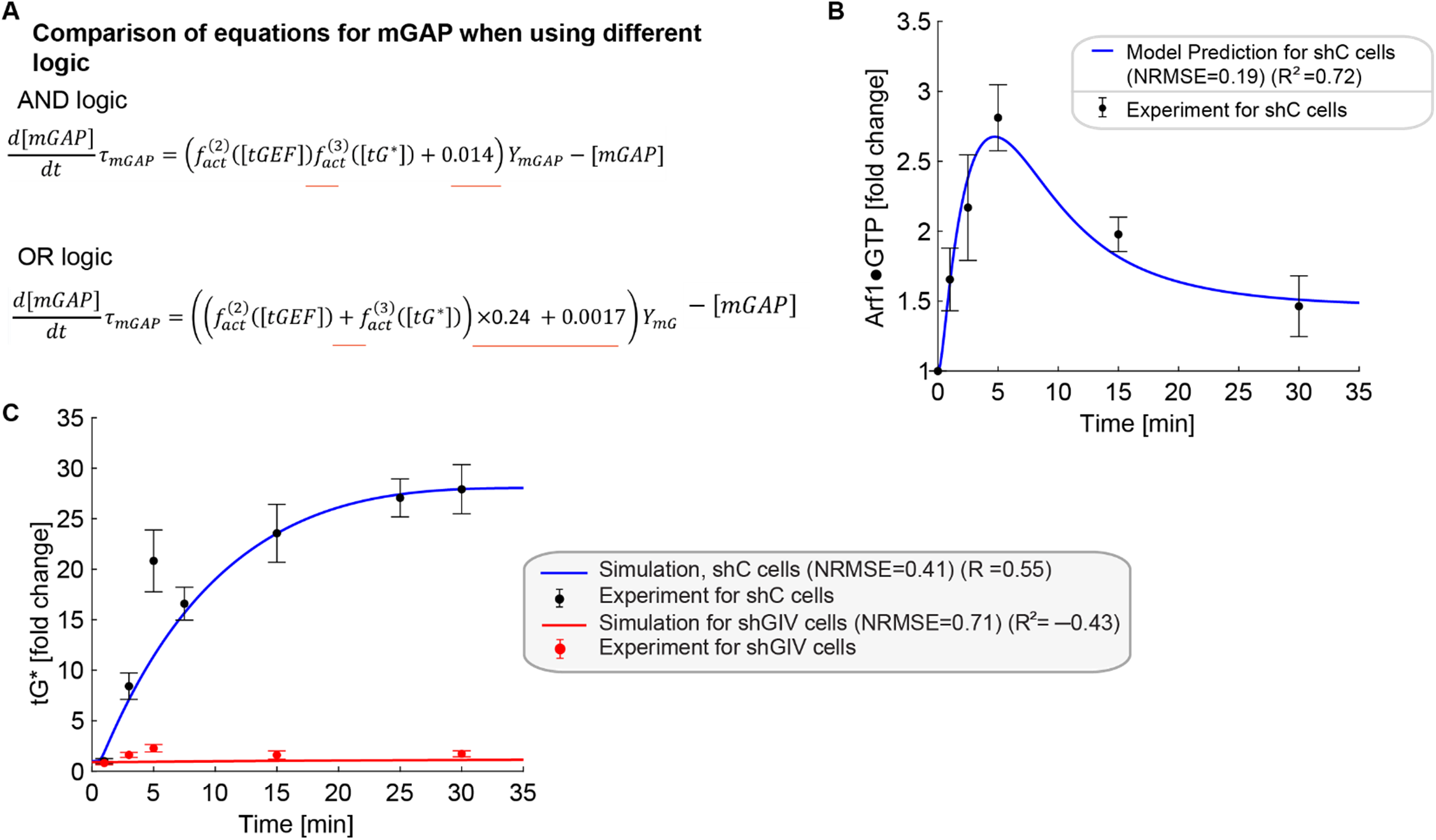
Simulations of Arf1 activation dynamics (mG*) and Gi activation dynamics (tG*) when using OR logic. **A.** Comparisons of equations for mGAP when using different logics. The AND and OR logic are modeled by 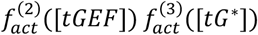 and 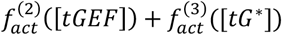, respectively. The constants 0.24 and 0.0017 ensure that the steadystate values of all species when using OR logic are the same as those for AND logic. The differences in the two models are underlined in red in the equations. **B-C.** Simulations of Arf1 (B) and Gi (C) activation dynamics for OR logic. The OR logic also captures the experimental data reasonably well but the AND logic is better informed experimentally.

**Figure EV4.**
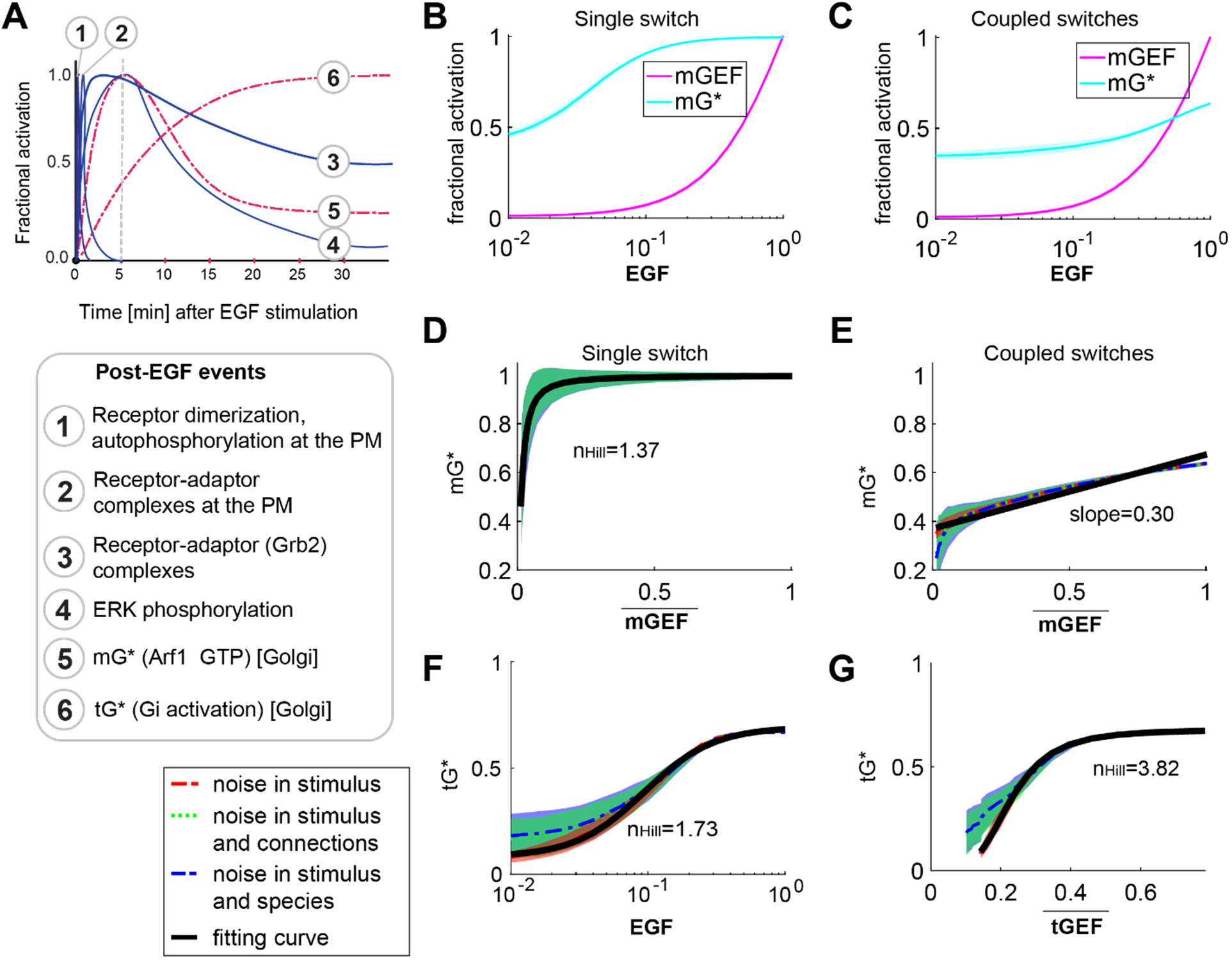
Coupled switches enable the alignment of endomembrane responses (Arf1 and tG* activities) to the dose of extracellular stimulus. **A.** Published dynamics of EGF-stimulated events that are initiated at the PM (blue, continuous) or experimentally determined dynamics of events at the Golgi confirmed here (red, interrupted) are compared. **B-C.** Dose responses of fractional activations of mGEF and active Arf1 (mG*) for the single switch (B; mG alone) and coupled switches (C; mG and tG). We perform stochastic simulations in the presence of noise in EGF (see **Materials and Methods** for details). The mean and the standard deviation (SD) of species are evaluated at steady states. The dimensionless EGF concentrations in the simulations are obtained through normalization, i.e., dividing the EGF concentration by 217.4 nM (=50 nM/0.23). In all simulations, noise is introduced only in stimulus (i.e., EGF). **D-E.** The same plots as in **Figure 4A-B** but in the presence of three different types of noise: noise in stimulus (shown in red), noise in stimulus and connections simultaneously (shown in green), and noise in stimulus and species simultaneously (shown in blue) (see **Materials and Methods** for details). Data are shown as mean values (dashed curves), with the shading showing the SD. The black curves are fitting curves (r^2^>0.95) for red dashed curves (see **Materials and Methods** for the calculations of r^2^ and *n_Hill_*). **F-G.** The fractional activations of tGTPase (tG*) as a function of EGF (F) or tGEF (G). The plots are generated in a similar way as in F-G. 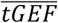 denotes the mean of tGEF. r^2^>0.95 for all fitting curves. The EGF-tG* relationship shows a Hill coefficient of 1.73, and the tGEF→tG* switch (switch #2) shows a dose-response behavior close to the saturation regime of an ultrasensitive curve (n_Hill_=3.82).

**Figure EV5.**
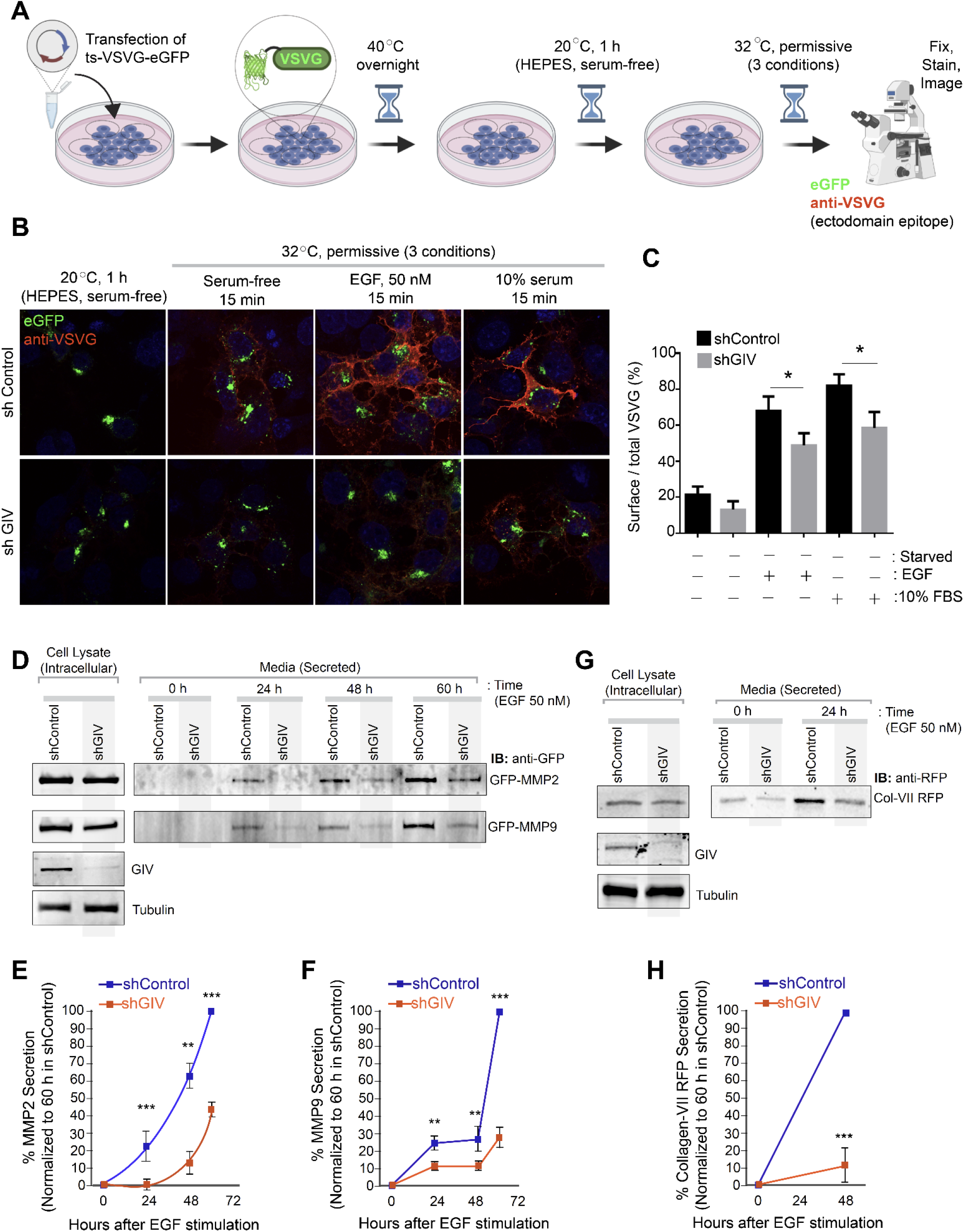
GIV-GEM is required for EGF-triggered secretion of diverse cargo proteins through Golgi compartment. **A**. Schematic shows the basis for measuring secretion of transmembrane cargo protein, ts-VSVG-eGFP. This temperature-sensitive mutant VSVG is retained in the ER at 40°C, at the Golgi at 20°C, and moves out of the Golgi to the PM when shifted to 32°C (Gallione and Rose, 1985). When visualized with immunofluorescence under non-permeabilized conditions, a VSVG-ectodomain targeting antibody selectively detects PM-localized cargo, whereas GFP tag allows the visualization of total VSVG in the cell. **B-C**. Control (sh Control; top) and GIV-depleted (shGIV; bottom) Cos7 cells were transfected with tsO45-VSVG-GFP and cells were shifted to 40°C for O/N and then incubated at 20°C for 1 h in HEPES buffered serum free media followed by temperature shift at 32°C for 15 minutes in plain DMEM and or containing 50nM EGF or 10% serum. Coverslips were fixed and stained with VSVG-ectodomain specific monoclonal antibody (red). Representative images are shown in B. Green fluorescence indicates total VSVG expression whereas red fluorescence shows surface-localized pool of VSVG. Bar graphs in C display the Red:Green intensity ratio indicative of fraction VSVG that is secreted to the cell surface. Results are expressed as mean ± S.E.M; n = 3 replicates; *p* values were determined using Mann-Whitney t-test compared to t0: *, <0.05. **D-H**. Control (sh Control) and GIV-depleted (shGIV) HeLa cells were analyzed for EGF-stimulated secretion of three soluble cargo proteins, MMP2 (D, E), MMP9 (D, F) and Collagen-Vii RFP (G, H), as detected from the supernatants at indicated time points after EGF stimulation. Results are expressed as mean ± S.E.M; n = 3 replicates; *p* values were determined using Mann-Whitney t-test compared to t0: *, <0.05; **, <0.01; ***, <0.001. Immunoblots are representative of findings from at least 3 independent repeats.

## MATERIALS AND METHODS

### REAGENTS AND TOOLS TABLE

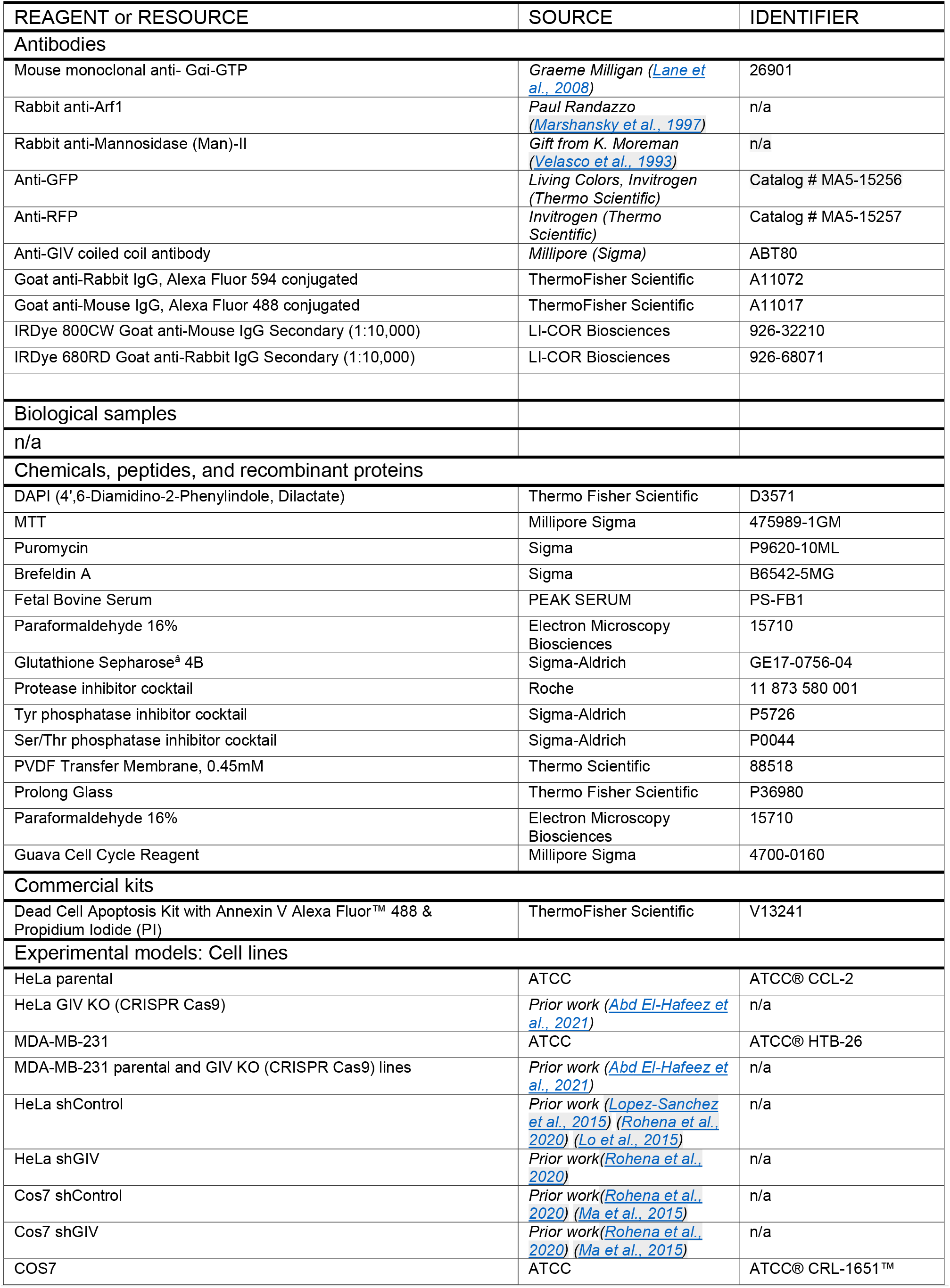
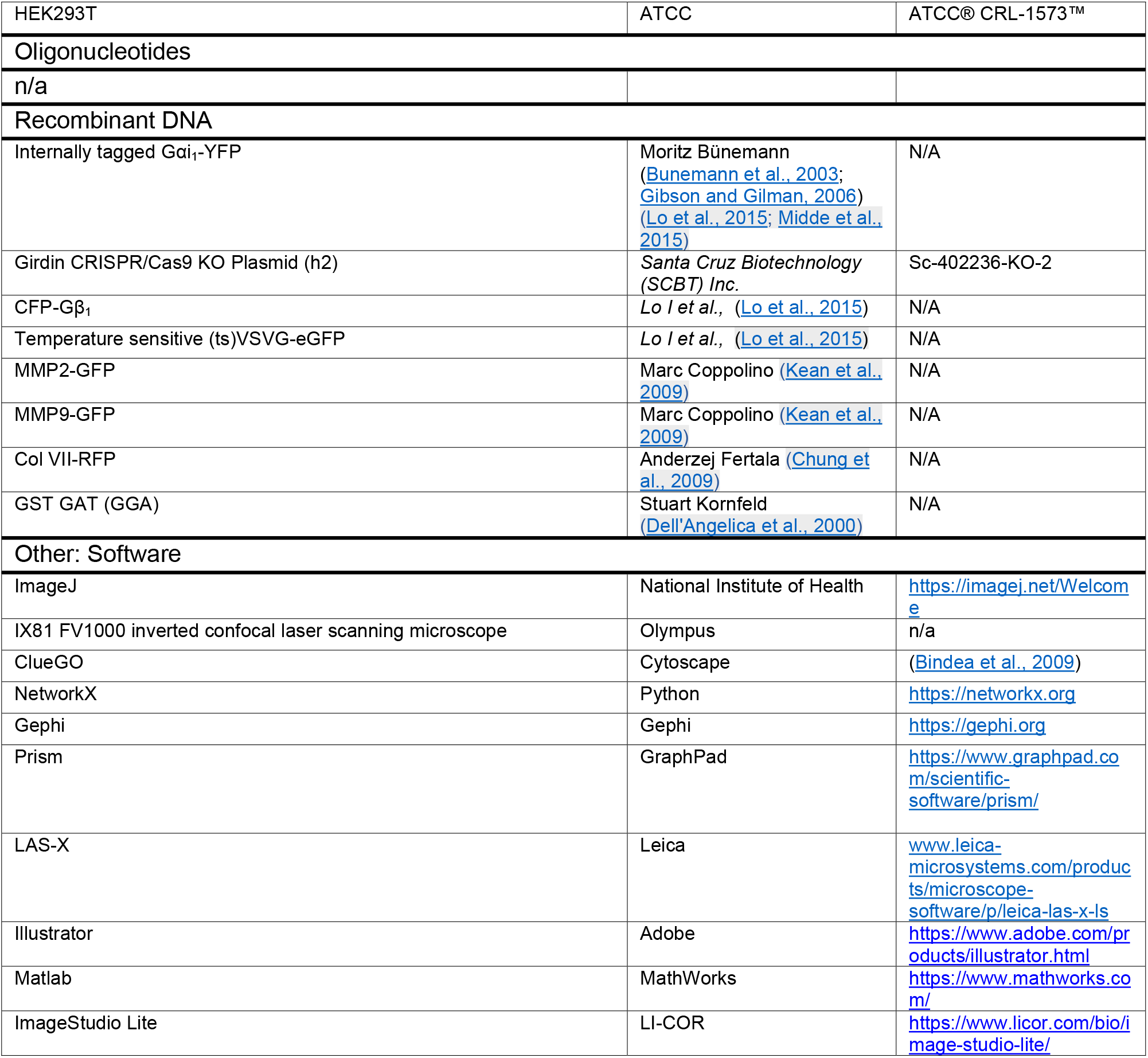

## METHODS AND PROTOCOLS

### Modeling approaches

#### Model Assumptions

We restrict our modeling considerations to the secretory pathway on Golgi and its interactions with the cell survival. The secretory pathway on Golgi consists of mGTPases, tGTPases, their GEFs and GAPs, and the secretion machinery. In the secretory pathway on Golgi, EGF mediates the recruitment of GEF for mGTPase (mGEF) and triggers the activation of corresponding mGTPases. Then active mGTPase can recruit GIV to vesicles. GIV is GEF for tGTPase (tGEF), and subsequently activates tGTPase. Upon activation of tGTPase, Gβγ is released and activates the GAP for mGTPase (mGAP). Besides, mGAP is also regulated by GIV, which binds to mGAP and works as a co-factor for GAP activity. mGAP has a dual role in this circuit: one is to turn ‘OFF’ mGTPase, and the other is to promote the vesicle formation. The vesicle formation is essential for secretion, and the secreted growth factors lead to the cell proliferation. The increase of cell number in turn enhances the secretion.

To model the above circuit, we assume that

- The total number of each type of GTPases is a constant.
- The copy number of GAP for tGTPase (tGAP) is constant since it is not regulated by other species.
- The species are present in large enough quantities that deterministic approaches can be used to capture the dynamics of the system.
- The process of secretion can be modeled using a simplified function that depends on mGAP.

Therefore, the circuit is modeled by a set of ordinary differential equations with six species: active mGTPase, active tGTPase, mGEF, mGAP, tGEF, and the secreted growth factors. Besides, the cells survival number is also modeled by an ordinary differential equation. We note that our model does not include the spatial or mechanical aspects associated with these signaling pathways.

#### Governing equations

Our model consists of two parts: one experimentally constrained module for coupled switches on the Golgi and the other module to predict the influence of coupled switches on the secrete-and-sense autonomy (**Figure 1B**). In the module for coupled switches, we modeled all the species interactions by normalized-Hill functions (Cao et al., 2020; Saucerman and McCulloch, 2004) to capture the overall input-output relationships. We did not consider all the intermediary steps in the signaling pathway for the sake of simplicity. When active tGTPase and tGEF both regulate mGAP, the “AND” logic is applied and modeled as *f_act_*(tGTPases) · *f_act_* (tGEF). Thus, the dynamics of the system can be described by the following equations:

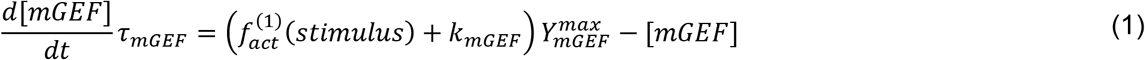

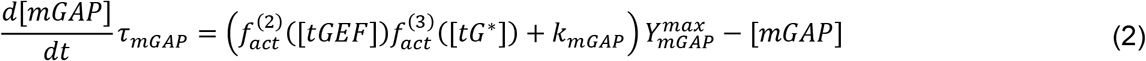

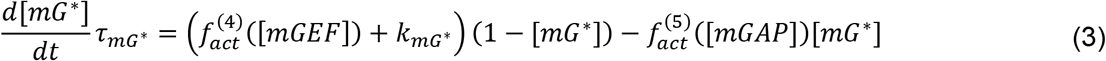

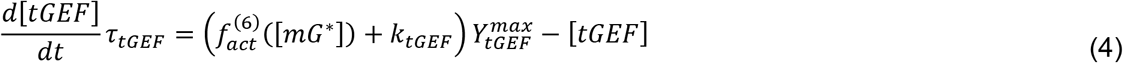

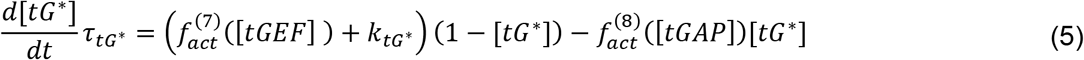

where variables [*mGEF*], [*mGAP*], [*mG*\*], [*tGEF*],and [*tG*\*] denote the fractional activation of mGEF, mGAP, mGTPase, tGEF, and tGTPase, respectively. Here, the fractional activation is the copy number divided by the maximal copy number, which changes between 0 and 1. The variable *stimulus* denotes the input signal EGF; the *τ*’s are time scale; *k*’s are basal production rates, and 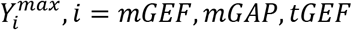 are maximal fractional activations for species. The function 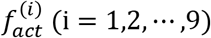 is the normalized-Hill function, which takes the following form:

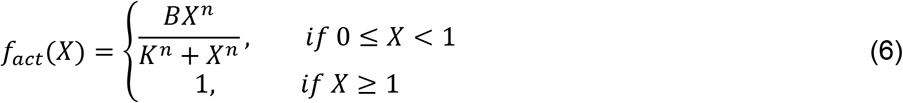

where 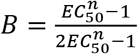 and *K* = (*B* – 1)^1/*n*^. Here, *EC*_50_ and *n* are half-maximal activation and Hill coefficient respectively. With these choices of constants *B* and *K*, we have *f_act_*(0) = 0, *f_act_*(*EC*_50_) = 0.5. It should be noted that constants *B* and *K* can be different in different functions 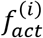. In most cases, we used *k* = 0 and *Y^max^* ≤ 1, so the maximal value of variables (1 + *k*)*Y^max^* is smaller than 1 to ensure the range of the fractional activation. But, when we used a non-zero *k*, the variable may be larger than 1, and then we regard the variable as the relative activation, which is normalized by a number smaller than the maximal copy number. We refer to this model as the coupled system throughout our study.

To predict the effect of coupled switches on the secrete-and-sense autonomy, we also built a model for secretion (denoted by S) and cell number (denoted by *X*). Since the activation-deactivation circle of mGTPase is necessary for the secretion, we assume the secretion rate is positively correlated to 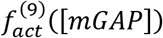. In addition, the proliferation of cells is regulated by secreted growth factors to ensure the homeostasis (Adler et al., 2018; Hart et al., 2014). Then, the dynamics of S and *X* are governed by:

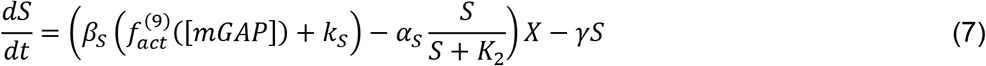

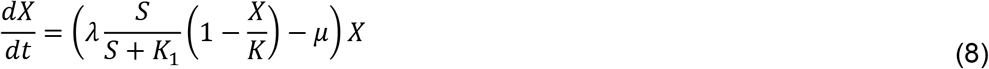

where *β_s_* is the maximal secretion rate; *k_s_* is the basal secretion rate; *α_s_* is the maximal endocytosis rate; *γ* is the degradation rate of secreted growth factors; *K*_2_ is the binding affinity of secreted growth factors. In the equation (8), *λ* and *μ* are cell proliferation and death rates by the cells, respectively; *K*_1_ is the value of S when the Hill function 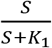 is 0.5; *K* is the carrying capacity, i.e., once the cell number is *K*, the cell proliferation rate is zero, preventing the cell number from exceeding *K*.

### Single switch model

For the circuit that only contains the single switch of mGTPases, its dynamics is described by equations (1)-(3), except that the equation (2) is replaced by

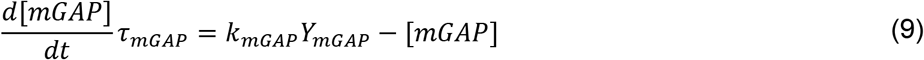

Note that this equation also can be used to describe the dynamics of mGAP when the regulation from tGEF to mGAP or the regulation from active tGTPase to mGAP doesn’t exist.

#### Numerical Simulations for the deterministic model

Numerical simulations were implemented in MATLAB. We use the solver ode15s to simulate the dynamics on the time interval [0,1440] minutes unless otherwise specified.

#### Fitting against experimental data

To fit the time course data for control cells and GIV-depleted cells, we manually tuned the parameters in our model until the normalized RMSE between simulated and measured fold changes of active Arf1 was less than 0.2 and that for active tGTPase less than 0.45. Moreover, parameters for the secretion and cell survival are taken from their biologically plausible ranges (Adler et al., 2018). Our fitting goal was to capture the experimentally observed trends rather than obtain kinetic parameters since our model does not include all the reactions in the pathway(s). Here, the normalized RMSE is the RMSE over the mean value of all experimental data; the baseline for the simulation result is the initial fractional activation when simulating dynamics for control cells, and those for experimental Arf1 and tGTPase data are initial states in control cells. The obtained parameter values are listed in **Table EV1**. In all simulations, the initial condition is the starved state when *stimulus* = 0, and then *stimulus* is set to be 0.23 to simulate the dynamics under the EGF-stimulated condition. In all simulations, we use normalized values of EGF concentrations. The normalization was conducted such that the value of 0.23 EGF used in simulations corresponds to 50 nM in the experiments. The dimensionless EGF concentrations in the simulations are obtained by dividing the EGF concentration by 217.4 nM (=50 nM/0.23).

#### Testing model

We verify that our setting in the model for GIV-deplete cells indeed makes the system behave like the uncoupled system. We set the maximal fractional activation of tGEF as 0.1 (i.e., *Y_tGEF_* = 0.1) but keep other parameters unchanged to model the system in GIV-depleted cells. The initial state is determined by the steady-state values of all species when the stimulus is zero, which are obtained as follows: we set *stimulus* zero and chose an arbitrary initial condition (e.g., all species are 0.5), and then simulated the deterministic dynamics on the time interval [0, 2400hr] to ensure that the steady state is reached. →Then we changed the stimulus to 0.23 to simulate the dynamics of all species when EGF=50 nM. We find that GIV-depleted cells more likely behave as the uncoupled system (**Appendix Figure S3A**). For these two systems, mGEF and mG* both increase upon the stimulus of EGF, and mGAP won’t increase because of low activation of tGEF in GIV-depleted cells or the absence of the positive regulation from tGEF and tG* in the uncoupled system. Due to non-increasing level of mGAP, these two systems both show non-decreasing fractional activation of mG*, low secretion and low cell number. The only difference between these two systems is the dynamics of tGTPase switch: tGEF is low in GIV-depleted cells and thus cannot activate tG*, while in the uncoupled system the fractional activations of tGEF and tG* are both high. The schematics of these three systems are shown in **Appendix Figure S3B-D**.

#### Sensitivity analysis

To test the robustness of the model, we performed the sensitivity analyses. The sensitivity measures how the system output is vulnerable to the parameter change, and can be captured by the following quantity:

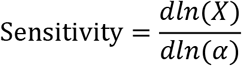

where the *X* is the system’s output, and *α* is the kinetic parameter. In our analyses, we calculated this sensitivity for each kinetic parameter, i.e., the *α* can be every kinetic parameter. The output *X* is the normalized RMSE value for simulated mG* or tG* dynamics, or steady-state values of the secretion or the cell number. This derivative is approximated by the ratio of the difference of *ln(X*) when *α* is 1.1 × *α* and 0.9 × *α* to the 0.2 × *α*. We found that, perturbations of the half-maximal activation *EC*_50_ will cause large changes for the normalized RMSEs and the steady-state value of the secretion for coupled switches (**Appendix Figure S4**). Except *EC*_50_, the mG* and tG* dynamics seem robust to other kinetic parameters, since the sensitivities for other kinetic parameters are between −0.5 and 0.5. Besides, the steady-state of the secretion in coupled switches is sensitive to the maximal secretion rate *α_s_* and the maximal endocytosis rate *β_s_*. These not very large sensitivities indicate that the main conclusions hold under small perturbations.

#### The stochastic model

To investigate the impact of noise, we consider three different sources of noise: stimulus, species and connections. A noisy stimulus is modeled by the summation of the mean and a noise term *η_sti_*(*t*); another type of noise, originated from species, is generated by adding a noise term *η_spe_*(*t*) in the equation for each species, and *tGAP* is also perturbed by a noise term 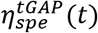; the third type of noise, which comes from connections, is modeled by adding a noise term *η_link_*(*t*) to each activation function *f_act_* and nonlinear reaction rates in equations for the secretion and the cell number. Here, these noise terms are independent from each other, and all modeled by the following Ornstein-Uhlenbeck process:

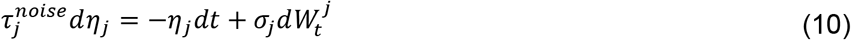

where *j* = *sti, spe, link*, and 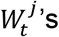 are independently and identical distributed standard Wiener processes. This equation implies that *η_j_*(*t*) has zero mean and variance 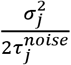. The equations for active tGEF, the secretory protein and the cell number in the presence of noise are taken as an example: when noise exists only in species, the dynamics of active tGEF, the secretory protein and the cell number are described by

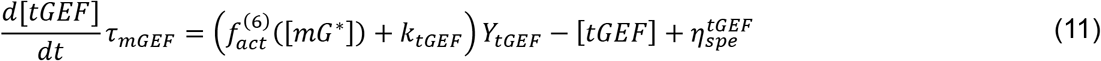

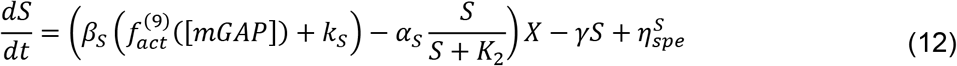

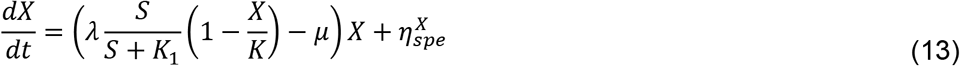

while the corresponding dynamics when noise are present in connections are governed by

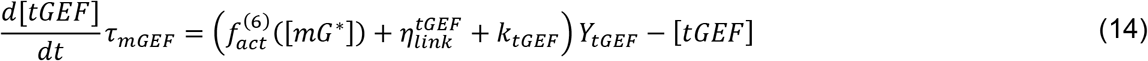

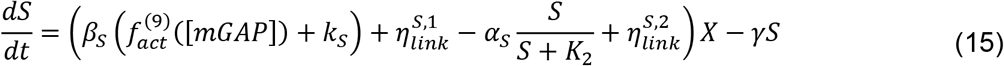

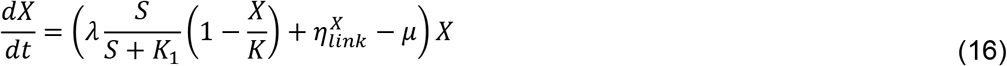

where *p_1ink_*’s with different superscripts are independent noise terms.

#### Numerical Simulations for the stochastic model

Numerical simulations were implemented in MATLAB. We used the Milstein scheme (Kloeden and Platen, 1992) to numerically solve the noise term *ty (j = sti, spe, link*), and used the Euler scheme to solve the dynamics of molecules on the time interval [0,1440min]. To be specific, the noise term *η_t_* at *n* +1 time step is determined by the following manner 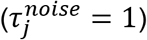:

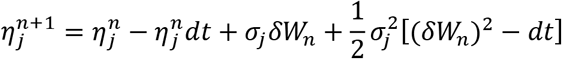

where *dt* is the time step, and *δW_n_* obeys the normal distribution with mean zero and variance *dt*. Then, the activation of molecules or the cell number is solved by the Euler scheme. For example, when noise is only in stimulus, the mGEF at *n* + 1 time step, denoted as [*mGEF*]^*n*+1^, is obtained by the following equation:

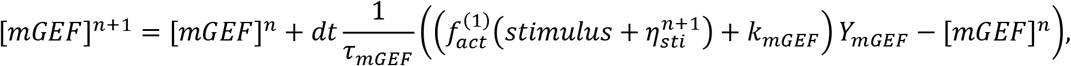

the schemes to solve the equations (12) and (15) are

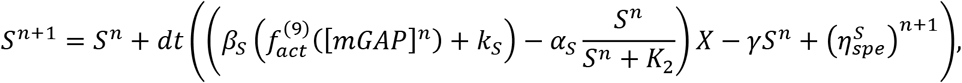

and

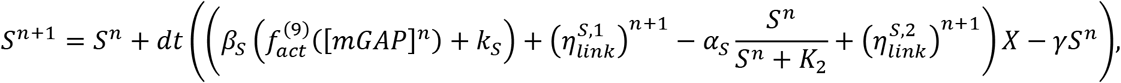

respectively.

We compare coupled switches with the single switch of mGTPase for three different cases of noise: noise in the stimulus, noise in the stimulus and species simultaneously, and noise in the stimulus and connections simultaneously. The values of noise amplitudes used for simulations are listed as follows:

- When noise is only in the stimulus, the parameter *σ_sti_* for *η_sti_*(*t*) is 0.02, and 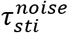 is 1.
- When noise is in the stimulus and species simultaneously, parameters *σ_sti_* and 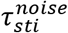 for *η_sti_*(*t*) are the same as those when noise is only in stimulus. In addition, for the noise term *η_spe_*(*t*), 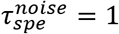, and *σ_spe_* is 0.02 for all species except the secretion and cell number. Since the secretion and cell number have small reaction rates, 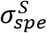 and 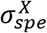 are set to be 2× 10^-5^ and 2 × 10^-6^ respectively, and thus the noisy behaviors cannot overwhelm the deterministic behaviors.
- When noise is in the stimulus and connections simultaneously, parameters *σ_sti_* and 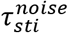 for *η_sti_*(*t*) are still the same as those when noise is only in stimulus. Moreover, 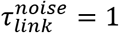, and 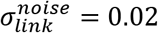 for all species except that the cell number. The 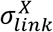 is 0.002 to ensure the same order of the noise and the production rate of cell number.

In this study, for a given input signal, we performed 1000 repeated simulations on the time interval [0 1440min] (with the steady state under this signal as the initial state). The time step *dt* is set to be 0.01.

### Computational and bioinformatics approaches

#### Identification of a Golgi-localized Arf1 and GIV interactome

We have previously extracted an annotated subcellular localization network of high confidence GIV correlators (Ear et al., 2021), based on Human Cell Map (HCM(Go et al., 2021)). From the same HCM data set, a set of high confidence Arf1 correlators were also extracted. Using the combined set of proteins that were correlated with GIV and Arf1, a full correlation network between every protein was extracted. Annotated unique GIV interactors from BioGRID (Oughtred et al., 2021) were also incorporated to expand the GIV-Arf1 interaction network. To assign subcellular localization of the GIV interactors from BioGRID (Oughtred et al., 2021), they were first matched to subcellular localization as annotated by HCM. For those proteins that were not assigned by HCM, they were then matched to Gene Ontology (GO) Cellular Component terms, Uniprot (The UniProt, 2019) and Human Protein Atlas (Uhlén et al., 2015), which were all used as a guide to manually assign them based on their biological function. The complete list of this ‘Golgi-localized Arf1-GIV interactome’ is provided as **Dataset EV1**.

#### Protein-protein interaction network construction, in silico perturbation and topological analyses

The list of proteins (**Dataset EV1**) was used as ‘seed’ to generate the Golgi-specific Arf1-GIV network by fetching other connecting interactions and proteins from STRING database (Franceschini et al., 2013). The shortest path NetworkX algorithm (Sinha et al., 2021) was used to trace the connected proteins and interactions in between every possible pair of protein from the above-mentioned list. Highest possible interaction cutoff score was used to avoid false positive interactions. To understand the impact of GIV deletion, a similar network was prepared, except without GIV. Shortest path alteration fraction (Sinha et al., 2021) associated with Arf1 was calculated using differential shortest path analysis of the original and GIV-depleted PPI network. Here only the paths having shortest path alteration fraction 1, were considered which indicated only the deleted or newly added shortest paths due to GIV deletion. GO Biological Process (BP) analysis of the proteins identified using shortest path alteration fraction analysis was performed using the Cytoscape tool ClueGO (Bindea et al., 2009) and significant GO BP terms were visualized.

#### TMT proteomics analysis, network construction and multi-layer visualization

Proteins that are upregulated in WT were mapped using the STRING database (https://string-db.org/). A pathway enrichment analysis of the most highly connected nodes was performed using the Reactome database (https://reactome.org/). The compartmentalized distribution of proteins within the PPI network based on their organelle-specific location was mapped using the Cell Atlas Uniform Manifold Approximation and Projection (UMAP) explorer that was generated using the large collection of confocal microscopy images showing the subcellular localization patterns of human proteins, curated and made available at Human protein atlas (https://www.proteinatlas.org/). Multilayer visualization of organelle-based interaction network was constructed using MultiViz plugin (S et al., 2022) of Gephi platform. All the source codes for network analysis are available at https://github.com/sinha7290/PPIN. MultiViz plugin source code available at https://github.com/JSiv/gephi-plugins.

### Experimental Model and Subject Details

#### Cell lines and culture methods

HeLa, Cos7 and MDA-MB-231 cells were grown at 37°C in their suitable media, according to their supplier instructions, supplemented with 10% FBS, 100 U/ml penicillin, 100 μg/ml streptomycin, 1% L-glutamine, and 5% CO2.

#### GIV CRISPR/Cas9 Gene Editing and Validation

Pooled guide RNA plasmids (commercially obtained from Santa Cruz Biotechnology; Cat# sc-402236-KO-2) were used to generate both HeLa and MDA MB-231 GIV KO lines as described before (Ear et al., 2021). Briefly, these CRISPR/Cas9 KO plasmids consist of GFP and Girdin-specific 20 nt guide RNA sequences derived from the GeCKO (v2) library and target human Girdin exons 6 and 7. Plasmids were transfected into Hela and MDA-MB-231 cells using PEI. Cells were sorted into individual wells using a cell sorter based on GFP expression. To identify cell clones harboring mutations in gene coding sequence, genomic DNA was extracted using 50 mM NaOH and boiling at 95°C for 60mins. After extraction, pH was neutralized by the addition of 10% volume 1.0 M Tris-pH 8.0. The crude genomic extract was then used in PCR reactions with primers flanking the targeted site. Amplicons were analyzed for insertions/deletions (indels) using a TBE-PAGE gel. Indel sequence was determined by cloning amplicons into a TOPO-TA cloning vector (Invitrogen) following manufacturer’s protocol.

#### Reagents and antibodies

All sources for key reagents are listed in the *Resource Table* above. Unless otherwise mentioned, all chemicals were purchased from Sigma (St Louis, MO). A mouse mAb against the active conformation of Gαi was obtained from Dr. Graeme Milligan (University of Glasgow, UK). Rabbit anti-Arf1 IgG was prepared as described (Marshansky et al., 1997). Rabbit polyclonal anti-α-mannosidase II (Man II) serum was prepared as described (Velasco et al., 1993). Highly cross-absorbed Alexa Fluor 594 or 488 F(ab)’2 fragments of goat anti-mouse or anti-rabbit IgG (H+L) for immunofluorescence were purchased from Invitrogen (Carlsbad, CA). Goat anti-rabbit and anti-mouse Alexa Fluor 680 or IRDye 800 F(ab)’2 for immunoblotting, were obtained from LI-COR Biosciences.

#### Cell culture, transfection, ligand stimulation and lysis

HeLa and MDA MB-231 (American Type Culture Collection, Manassas, VA) were maintained in DMEM (Invitrogen) supplemented with 10% FBS (Hyclone, Logan, UT), 100 U/ml penicillin, 100 μg/ml streptomycin, 1% L-glutamine and 5% CO2. Control and GIV shRNA HeLa and Cos7 stable cell lines were selected with 2 μg/ml of Puromycin (GIBCO) using plasmid expressing a shRNA targeting its 3’ UTR (Ghosh et al., 2016a). Depletion of GIV was verified using GIV-CT antibody with an efficiency of ~95% and cells were extensively validated in prior studies (Lo et al., 2015; Ma et al., 2015; Rohena et al., 2020). Transfection of cells with fluorescent plasmids (FRET studies) was carried out using transit-LT1 (Mirus Bio, Madison, WI) following the manufacturer’s protocol. Cells were checked for mycoplasma contamination and authenticated by STR profiling periodically.

For ligand stimulation of cells, serum starvation was carried out overnight (~16-18 h) by replacing media with 0.2% FBS containing media in the case of HeLa prior to exposing them to the ligands.

Lysates used as a source of proteins in pulldown assays were prepared by resuspending cells in Tx-100 lysis buffer [20 mM HEPES, pH 7.2, 5 mM Mg-acetate, 125 mM K-acetate, 0.4% Triton X-100, 1 mM DTT, supplemented with sodium orthovanadate (500 mM), phosphatase (Sigma) and protease (Roche) inhibitor cocktails], after which they were passed through a 28G needle at 4°C, and cleared (10,000 x g for 10 min) before use in subsequent experiments.

#### Arf1 activation assays

Purification of GST-GAT protein and assessment of Arf1 activation was described previously. In brief, cells were lysed with 1% Triton X-100, 50 mM Tris, pH 7.5, 100 mM NaCl, 2 mM MgCl2, 0.1% SDS, 0.5% sodium deoxycholate, 10% glycerol with protease inhibitors. Equal amounts of lysates were incubated with GST-GGA3 (~40 μg) prebound glutathione-Sepharose 4B beads at 4°C for 1 h. Beads were washed, and the bound proteins were eluted by boiling in Laemmli sample buffer for 5 min, resolved on a 15% SDS-PAGE, and analyzed by immunoblotting.

#### Quantitative immunoblotting

For immunoblotting, protein samples were boiled in Laemmli sample buffer, separated by SDS-PAGE and transferred onto 0.45 mM PVDF membrane (Millipore) prior to blotting. The duration of transfer was 30 min, at 100 V. Post transfer, membranes were blocked using 5% Non-fat milk or 5% BSA dissolved in PBS. Primary antibodies were prepared in blocking buffer containing 0.1% Tween-20 and incubated with blots, rocking overnight at 4*°*C. After incubation, blots were incubated with secondary antibodies for one hour at room temperature, washed, and imaged using a dual-color Li-Cor Odyssey imaging system.

#### Immunofluorescence and confocal microscopy

For immunofluorescence, cells grown on coverslips were fixed in 3% paraformaldehyde (PFA) and processed as described previously (Ghosh et al., 2010). Antibody dilutions were as follows: Man II, 1:800; anti-Gαi•GTP, 1:25; goat anti-mouse or anti-rabbit Alexa 488 or Alexa 594, 1:500. DAPI was used at 1:10,000. To estimate the degree of colocalization (Mander’s overlap coefficient; MOC) in immunofluorescence assays, an ImageJ plugin, JACoP (https://imagej.nih.gov/ij/plugins/track/jacop2.html) was used. This was preferred over Pearson’s because it is a good indicator of the proportion of the green signal (active G protein) coincident with a signal in the red channel (Man II, indicative of Golgi membranes) over its total intensity, which may even apply if the intensities in both channels are really different from one another. Coverslips were mounted using Prolong Gold (Invitrogen) and imaged using a Leica SPE CTR4000 confocal microscope.

#### Image processing

All images were processed on ImageJ software (NIH) and assembled into figure panels using Photoshop and Illustrator (Adobe Creative Cloud). All graphs were generated using GraphPad Prism.

#### Förster Resonance Energy Transfer (FRET) studies

Intramolecular FRET was detected by sensitized emission using the three-cube method were performed as previously reported by Midde et al (Midde et al., 2015). Briefly, previously validated internally tagged Gαi1-YFP and CFP-Gβ1 FRET probe pairs were used (Bunemann et al., 2003; Gibson and Gilman, 2006). Cells were transfected with the probes, serum starved overnight, and then stimulated with EGF (50 nM) exactly as done previously (Kalogriopoulos et al., 2020; Midde et al., 2015). All fluorescence microscopy assays were performed on single cells in mesoscopic regime to avoid inhomogeneities from samples as shown previously by Midde et al (Borejdo et al., 2012; Midde et al., 2015). Briefly, cells were sparsely split into sterile 35 mm MatTek glass bottom dishes and transfected with 1 μg of indicated constructs. To optimize the signal-to-noise ratio in FRET imaging, various expression levels of the transfected FRET probes were tested. However, to minimize complexities arising from molecular crowding, FRET probes were overexpressed by ~1.5- to twofold compared with the endogenous proteins. Because the stoichiometry of FRET probes has a significant impact on FRET efficiency, cells that expressed equimolar amounts of donor and acceptor probes (as determined by computing the intensity of the fluorescence signal by a photon-counting histogram) were chosen selectively for FRET analyses. An Olympus IX81 FV1000 inverted confocal laser scanning microscope was used for live cell FRET imaging (UCSD-Neuroscience core facility). The microscope is stabilized on a vibration proof platform, caged in temperature controlled (37°C) and CO2 (5%) supplemented chamber. A PlanApo 60x 1.40 N.A. oil immersed objective designed to minimize chromatic aberration and enhance resolution for 405-605 nm imaging was used. Olympus Fluoview inbuilt software was used for data acquisition. A 515 nm Argon-ion laser was used to excite EYFP and a 405 nm laser diode was used to excite ECFP as detailed by Claire Brown’s group (Broussard et al., 2013). Spectral bleed-through coefficients were determined through FRET-imaging of donor-only and acceptor-only samples (i.e. cells expressing a single donor or acceptor FP). Enhanced CFP emission was collected from 425-500 nm and EYFP emission was collected through 535-600 nm and passed through a 50 nm confocal pinhole before being sent to a photomultiplier tube to reject out of plane focused light. Every field of view (FOV) is imaged sequentially through ECFPex/ECFPem, ECFPex/EYFPem and EYFPex/EYFPem (3 excitation and emission combinations) and saved as donor, FRET and acceptor image files through an inbuilt wizard. To obtain the FRET images and efficiency of energy transfer values a RiFRET plugin in Image J software was used (Roszik et al., 2009). Prior to FRET calculations, all images were first corrected for uneven illumination, registered, and background subtracted. For FRET quantification, regions of interest (ROI) were drawn in the juxtanuclear area presumably in the Golgi region (or at the cell periphery, presumed to be the plasma membrane regions) to compute energy transfer. Individual cells with fluorescence intensity in the mesoscopic regime detected in the donor and acceptor channels were selected for FRET analysis to avoid inhomogeneities between samples (Midde et al., 2013; Midde et al., 2014)

Manual and automatic registration of each individual channel in ImageJ was critical to correct for motion artifacts associated with live cell imaging. Controls were performed in which images were obtained in different orders. The order in which images were obtained had no effect. FRET images were obtained by pixel-by-pixel ratiometric intensity method and efficiency of transfer was calculated by the ratio of intensity in transfer channel to the quenched (corrected) intensity in the donor channel. The following corrections were applied to all FOVs imaged: For crosstalk correction, cells transfected with CFP or YFP alone were imaged under all three previously mentioned excitation and emission combinations. FRET efficiency was quantified from 3-4 Regions of Interests (ROI) per cell drawn exclusively along the P.M. Because expression of FRET probes may have a significant impact on FRET efficiency, cells that expressed similar amounts of probes, as determined by computing the fluorescence signal/intensity by a photon counting histogram were selectively chosen for FRET analyses. Furthermore, untransfected cells and a field of view with-out cells were imaged to correct for background, autofluorescence and light scattering. To avoid artifacts of photobleaching, Oxyfluor (www.oxyrase.com) was used to minimize the formation of reactive oxygen species.

#### GFP-tsO45-VSVG transport assays

To monitor anterograde (ER to Golgi) trafficking control or GIV-depleted COS7 cells were transiently transfected with GFP-tsO45-VSV-G plasmid (Presley et al., 1997). Transfected cells were incubated for 14–16 h at the restrictive temperature (40°C) to accumulate VSV-G protein in the ER, shifted to 32°C for 0-60 min to release VSV-G protein in the conditions described (i.e., 10% serum, EGF, or starved condition) and then fixed and processed under non-permeabilized conditions (without detergent) for immunofluorescence. The rate of VSV-G trafficking from the secretory compartments to the PM was determined by calculating the ratio of VSV-G that was already at the PM (as determined using an anti-VSV-G ectodomain antibody; red pixels) normalized to the total cellular pool of VSV-G (GFP; green pixels, using NIH *ImageJ* software.

#### Metalloprotease and collagen secretion assays

HeLa cells grown in a 6 well plate were transfected with 2 ug of GFP-MMP2, GFP-MMP9 or Collagen-RFP for 5h. After 5 h, cells were fed with fresh media without FBS. Media was subsequently changed the next day (without FBS; exactly 1.5 ml/well) and stimulated with EGF. Media (100 ul) was collected just before the addition of EGF, as T= 0 h, and at the indicated time points after EGF stimulation. Each aliquot was subjected to high speed (14,000 x g) spin for 10 min prior to the addition of 50 ul of Laemmli sample buffer and boiling at 100 °C.

#### MTT assay

Cell proliferation was measured using the MTT reagent and cells cultured in 96-well plates. Parental or GIV-KO HeLaor MDA-MB-231 cells were cultured in different concentrations of Fetal Bovine Serum (FBS; 0, 0.25, 2, 5, and 10 %)Then the cell lines were incubated with MTT for 4 hr at 37°C. After incubation, culture media was removed and 150 μl of DMSO was added in order to solubilize the MTT formazan crystals. Optical density was determined at 590 nm using a TECAN plate reader. At least three independent experiments were performed. In an independent experiment we tested the effect of using a Brefeldin A (BFA), a well-known tool to inhibit secretion, on the cell proliferation. The cell lines were cultured in different concentrations of FBS (0, 0.25, 2, 5, or 10 %) and then treated with different concentrations of BFA (0, 0.01, 0.05, 0.1, 0.5, 1, 10, or 100 μM) and the MTT assays were done as described.

#### Cell cycle and apoptosis analyses

Cell cycle analysis and apoptotic cell quantification were performed using the Guava cell cycle reagent (Millipore Sigma) or the annexin V/propidium iodide (PI) staining kit (Thermo Fisher Scientific), respectively, according to the manufacturer’s instructions. Cells were quantified on a BD™ (BD Biosciences) LSR II flow cytometer and analyzed using FlowJo software (FlowJo, Ashland, OR, USA).

#### Tandem Mass Tag™ (TMT) proteomics

WT and GIV-KO MDA-MB231 cells were maintained in 0% and 10% serum concentration in p10 dishes (Corning) for 16 h prior to harvest, and cell pellets were subsequently processed for TMT proteomics using LUMOS Orbitrap-Fusion analyzer. Samples were processed at the UC San Diego Biomolecular and Proteomics Mass Spectrometry Core Facility (https://bpmsf.ucsd.edu/). Peptides are identified and mapped using Peaks X Pro pipeline. Intensity ratio of each identified proteins in WT MDA-MB231 Vs GIV-KO MDA-MB231 cells has been identified and selected if the significance score >20. A list of differentially expressed proteins is provided in **Dataset EV2**. The mass spectrometry proteomics data have been deposited to the ProteomeXchange Consortium via the PRIDE partner repository (Perez-Riverol et al., 2022) with the dataset identifier PXD037253.

## QUANTIFICATION AND STATISTICAL ANALYSIS

### Statistical analyses in modeling approaches

In the deterministic model, we fitted the dose-response curve by finding the best-fit function with the form 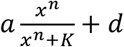. We solved this optimal problem using ‘lsqcurvefit’ in Matlab, and *d* can be deleted depending on the effect of the fitting. The only exception is for the mG versus mGEF, where we used linear function *αx* + *d*. The difference between the fitted curve and the original curve is measured by R^2^, and it is defined as 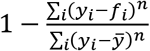, where *y_i_* is the point in the original curve and *f_i_* is the prediction for *y_i_* based on the best-fit curve. In the stochastic model, the standard deviation is calculated based on the data at 1440min, which is defined as the

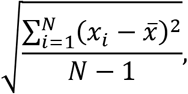

where *N* = 1000 and 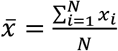. The *x_f_* is the molecular activation or the cell number at 1440min in the *i*-th simulation.

### Statistical analyses in protein-protein network analyses

An interaction cutoff score has been optimized while fetching the new proteins and their interactions from STRING database, such that all the possible proteins will be included keeping the cutoff very high. In this instance, an interaction cutoff score of 667 has been used to include all the proteins from the seed list (**Dataset EV1**).

### Statistical analyses in experimental studies and replication

All experiments were repeated at least three times (biological replicates, conducted on different days), and results were presented either as one representative experiment or as average ± S.E.M. Statistical significance was assessed with two-sided unpaired Student’s t test and Mann-Whitney t-test. For all tests, a p-value of 0.05 was used as the cutoff to determine significance. The actual p-values are indicated in each figure. All statistical analysis was performed using GraphPad prism 8 or Matlab. Experiments undertaken did not require blinding; nor did they require sample size calculation or randomization.

## DATA AVAILABILITY

- Modeling computer scripts: GitHub (https://github.com/RangamaniLabUCSD/Coupled-switches-secretion)
- Protein-protein network analyses: Github (https://github.com/RangamaniLabUCSD/Coupled-switches-secretion)
- TMT proteomics datasets: PXD037253 (http://www.proteomexchange.org)

## RESOURCE AVAILABILITY

### Lead Contact

Further information and requests for resources and reagents should be directed to and will be fulfilled by the Lead Contact, Pradipta Ghosh. All model-related queries could be alternatively directed to and will be fulfilled by Padmini Rangamani.

### Materials Availability

- This study did not generate new unique reagent

