## Supplementary material for "A Circuit for Secretion-coupled Cellular Autonomy in Multicellular Eukaryotes": Movie EV1: README.txt

Movie EV1. Dynamics within the endomembrane GTPase system. A movie gif that captures the dynamic nature of a circuit of two species of coupled GTPases, mGTPases (mG) and heterotrimeric GTPases (tG), that is localized on membranes within the secretory pathway. 
