## Supplementary figures and images for "A Circuit for Secretion-coupled Cellular Autonomy in Multicellular Eukaryotes"

### Movie EV1.gif

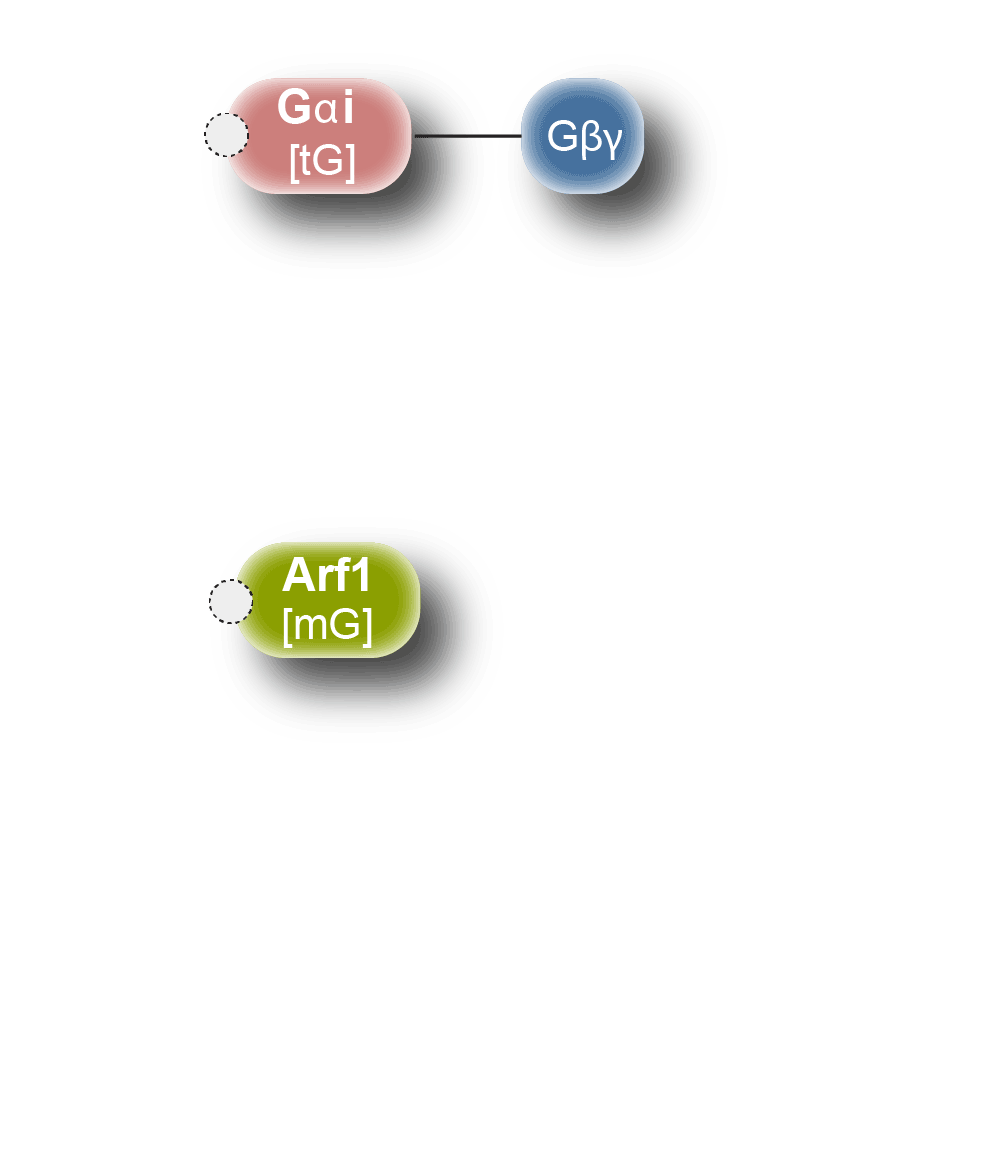
