## Supplementary material for "A Circuit for Secretion-coupled Cellular Autonomy in Multicellular Eukaryotes": Dataset EV2: README.txt

Dataset EV2. Excel sheet displays the list of proteins differentially expressed in coupled vs uncoupled HeLa and MDA MB-231 cells that were used to create the protein-protein interaction network in Figure 4.
