## Supplementary material for "A Circuit for Secretion-coupled Cellular Autonomy in Multicellular Eukaryotes": Dataset EV1: README.txt

Dataset EV1. Excel sheet displays the list of proteins identified as part of the Golgi-localized Arf1-GIV interactome and was used as ‘seed’ list for creating the Golgi localized protein-protein interaction network. 
